# Genetic testing of dogs predicts problem behaviors in clinical and nonclinical samples

**DOI:** 10.1101/2020.08.13.249805

**Authors:** Isain Zapata, M. Leanne Lilly, Meghan E. Herron, James A. Serpell, Carlos E. Alvarez

## Abstract

Very little is known about the etiology of personality and psychiatric disorders. Because the core neurobiology of many such traits is evolutionarily conserved, dogs present a powerful model. We previously reported genome scans of breed averages of ten traits related to fear, anxiety, aggression and social behavior in multiple cohorts of pedigree dogs. As a second phase of that discovery, here we tested the ability of markers at 13 of those loci to predict canine behavior in a community sample of 397 pedigree and mixed-breed dogs with individual-level genotype and phenotype data. We found support for all markers and loci. By including 122 dogs with veterinary behavioral diagnoses in our cohort, we were able to identify eight loci associated with those diagnoses. Logistic regression models showed subsets of those loci could predict behavioral diagnoses. We corroborated our previous findings that small body size is associated with many problem behaviors and large body size is associated with increased trainability. Children in the home were associated with anxiety traits; illness and other animals in the home with coprophagia; working-dog status with increased energy and separation-related problems; and competitive dogs with increased aggression directed at familiar dogs, but reduced fear directed at humans and unfamiliar dogs. Compared to other dogs, Pit Bull-type dogs were not defined by a set of our markers and were not more aggressive; but they were strongly associated with pulling on the leash. Using severity-threshold models, Pit Bull-type dogs showed reduced risk of owner-directed aggression (75^th^ quantile) and increased risk of dog-directed fear (95^th^ quantile). Our findings have broad utility, including for clinical and breeding purposes, but we caution that thorough understanding is necessary for their interpretation and use.

## Introduction

There are an estimated 70-90M pet dogs in 36.5-42% of US homes^1,2^. Because dogs suffer from many of the same conditions as humans and often receive a high level of health care, they represent an ideal comparative and translational animal model^3,4^. Strong positive selection in dog domestication and breed development had the by-effect of vastly relaxed negative selection. As a result, most complex traits studied to date in dogs – including cancer and other diseases, morphology and behavior – have shown dramatically-reduced polygenicity and moderate-to-large effect sizes of associated variants^3,5-13^. In contrast, common human complex disease variations generally have small effect sizes and are not directly useful medically or experimentally.

We previously reported genome wide association studies (GWASs) of breed averages for fear and aggression behaviors in multiple cohorts with different breed make-ups^11,12^. That was based on behavioral data using the validated survey instrument C-BARQ^14-16^. MacLean et al. performed similar scans of those and other behaviors in one shared and one different cohort, but correcting for body mass in the association analysis^13^. In a study partnered with the present work, we compared the scans of diverse behaviors with and without correction for body mass. The correlation of body size and behavior in dogs has thus been observed in behavioral^17-20^ and genetic studies^11,12^. Based on biological relevance, we previously argued that behavior-body size correlations are due to pleiotropy^12^ (rather than population structure^21^). We and others have now strengthened that evidence greatly^11,22^.

In much of the world, dogs exist as pedigree (stratified by hundreds of breeds), mixed-breed and free-roaming subpopulations (Suppl. Text). Pit Bulls, which are among the most popular dog types in the US, are a special and controversial case (Suppl. Text). The ancestral UK Staffordshire Bull Terrier was once selected for dog fighting and it is unclear to what degree that has continued or whether the breed type should be considered especially dangerous. Pit Bull refers to a group of breeds, some of which are registered by the American Kennel Club (AKC) and others by different institutions in the US^23^. A recent study of two dog shelters in different US states compared visual and genetic classification of Pit Bull-type dogs^24^. Shelter staff had a 76% correct call rate for 114 dogs that were genetically greater than 25% American Staffordshire Terrier (AST). Their false positive rate for 270 non-AST dogs was 1.5%. Of the total 919 dogs from both shelters, 238 had an AST genetic signature (24% and 28%) and the average AST contents were 39% and 48%. Below 25-38% AST content, the correct visual calling rate of Pit Bull-type dogs falls rapidly. A C-BARQ-based behavioral study of ∼3,800 AKC registered dogs from 32 breeds also included 132 Pit Bull-type dogs as defined here^17^. Pit Bull-type dogs showed significantly decreased aggression to owners, but increased aggression to dogs (they did not rank highest in any behavior). Here we mitigated visual calling of Pit Bull-type dogs by performing principal components analyses (PCA) with the set genetic markers genotyped. With these caveats, this is the first genetic study of Pit Bull behavior.

As clinical and lay access to genetic testing continues to accelerate rapidly, it is important to understand its utility. In order for genetic tests to be clinically actionable, they have to be useful in the observation, diagnosis or treatment of patients. Knowledge of increased genetic risk can indicate therapeutic intervention, initiation and interpretation of disease screening, and life planning^25^. In domesticated animals, the latter includes management of commercial/production traits, welfare and reproductive planning. Because complex traits in domesticated species often have greatly reduced polygenicity and increased effect sizes of variations compared to humans, the utility of genetic testing in veterinary medicine and animal sciences is greatly simplified. Our long-term goal is the development and validation of genetic testing for use by veterinary behaviorists as well as dog breeders, shelter administrators and owners.

The objective of this work was to provide further evidence for the previous interbreed findings in a community sample. Whereas our GWASs were performed using breed averages of C-BARQ traits and unrelated genotyped-cohorts with varied breed makeups, here we used individual-level C-BARQ phenotypes and genotypes for 20 markers at 13 behavioral GWA loci. Our 400-dog cohort included a typical proportion of pedigree and mixed-breed dogs for the US, and was representative of the community and the veterinary behavioral clinic. Only variations common across breeds could have been mapped with our approach and such variations are enriched for admixture^26^. That is consistent with our present results because correlations between unlinked markers (associated with population structure) and between behaviors were distinct. Our findings lend support for the genome scans and utility of genetic testing, but, in the Discussion, we advise caution on direct-to-consumer tests.

## Results

### Study design, cohort and genotyping

Previous GWA discovery was performed using breed averages of behavior and unrelated genotyped cohorts of diverse pedigree dogs. In contrast, the present study i) targeted a subset of those GWA loci; ii) used both pedigree and mixed breed dogs; iii) used dogs with individual-level phenotypes and genotypes; and iv) included the original behavioral traits and additional ones. Factoring the complexities of the quantitative and population genetics, and our power, this work is a second phase of discovery – with the GWASs being the first.

We designed our study to evaluate the performance of genetic markers as predictors of canine problematic behavior in the community. We recruited subjects without breed or geographical restrictions (Suppl. Text). Dog clinical background and demographic data were provided by owners in the form of paper questionnaires, while behavioral information was collected via electronic questionnaires (C-BARQ). Paper questionnaire and genotype data were considered predictor variables and C-BARQ traits were considered response variables.

Our dog cohort included a total of 397 dog subjects. Descriptive statistics of our sample are provided in Table 1. Our sample had an almost even female to male ratio (47:52%) and most were neutered (365 vs. 31). All dogs were considered pets, 16 were classified as working-purposed and 17 as competition-purposed (2 as both). 45% of our dogs were members of 77 pedigree or designer breeds (Suppl. Table S1) and 55% were mixed breed. This is similar to the US proportion of mixed-breed dogs of 51-53%^1,27^. Owners were asked to describe the breed make-up of their dog. We evaluated popularities of breeds in the US and in select US cities, and determined our cohort to be representative of a typical US community despite being geographically biased for Ohio (Suppl. Table S2). Owners most commonly acquired dogs in our study from shelters, breeders and acquaintances. Other sources were pet stores and rescue organizations. Many dogs previously had other owners (e.g., most shelter dogs). Our cohort was intentionally biased to have increased representation of dogs with a behavioral diagnosis: 30% of our sample, of which 21% of those (or 6.5% of all dogs) were under medication for problem behavior. 30% of our subjects had a non-behavioral medical condition. Lastly, we noted whether dogs lived with other dogs, animals or children.

**Table 1.**
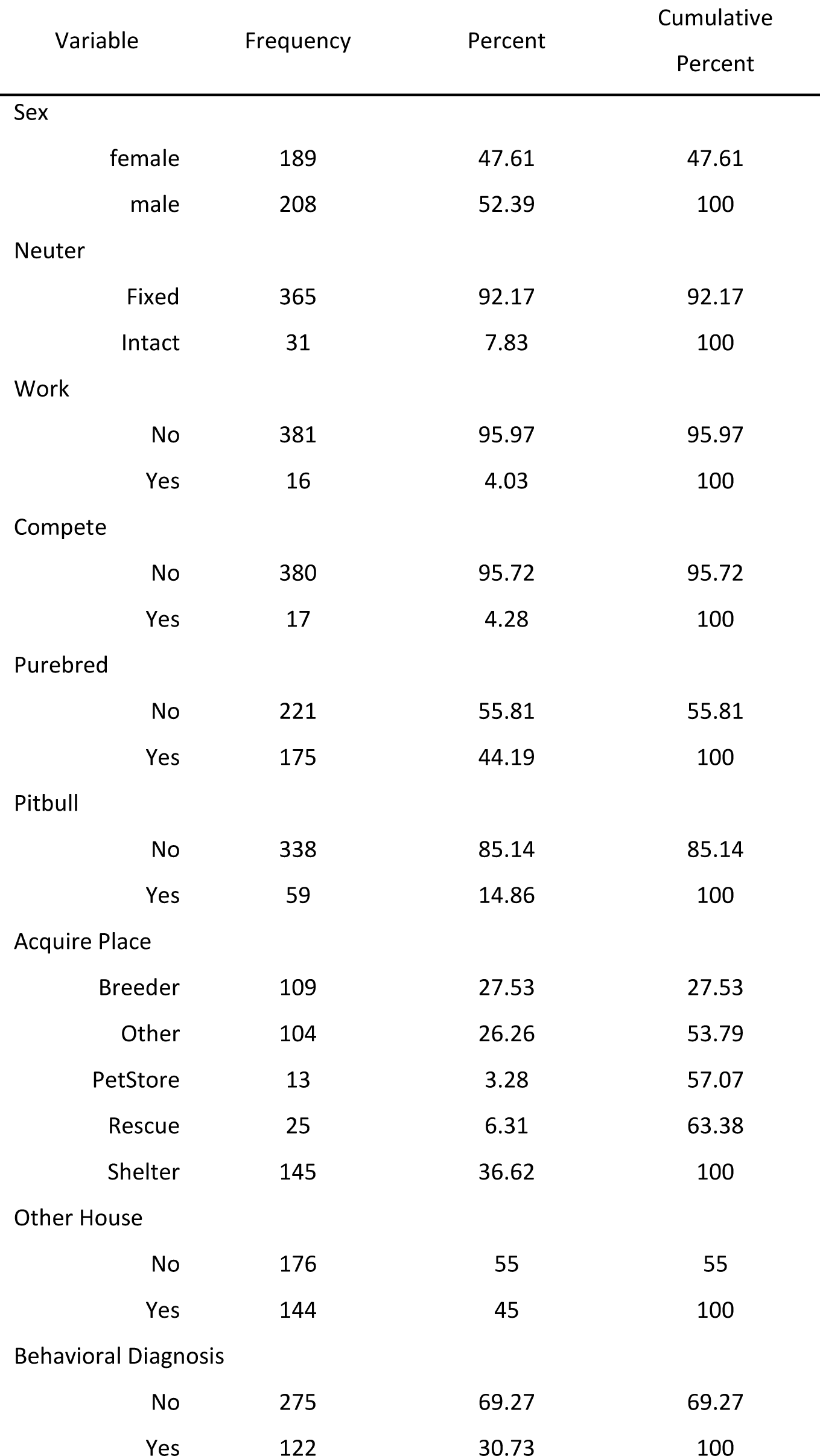
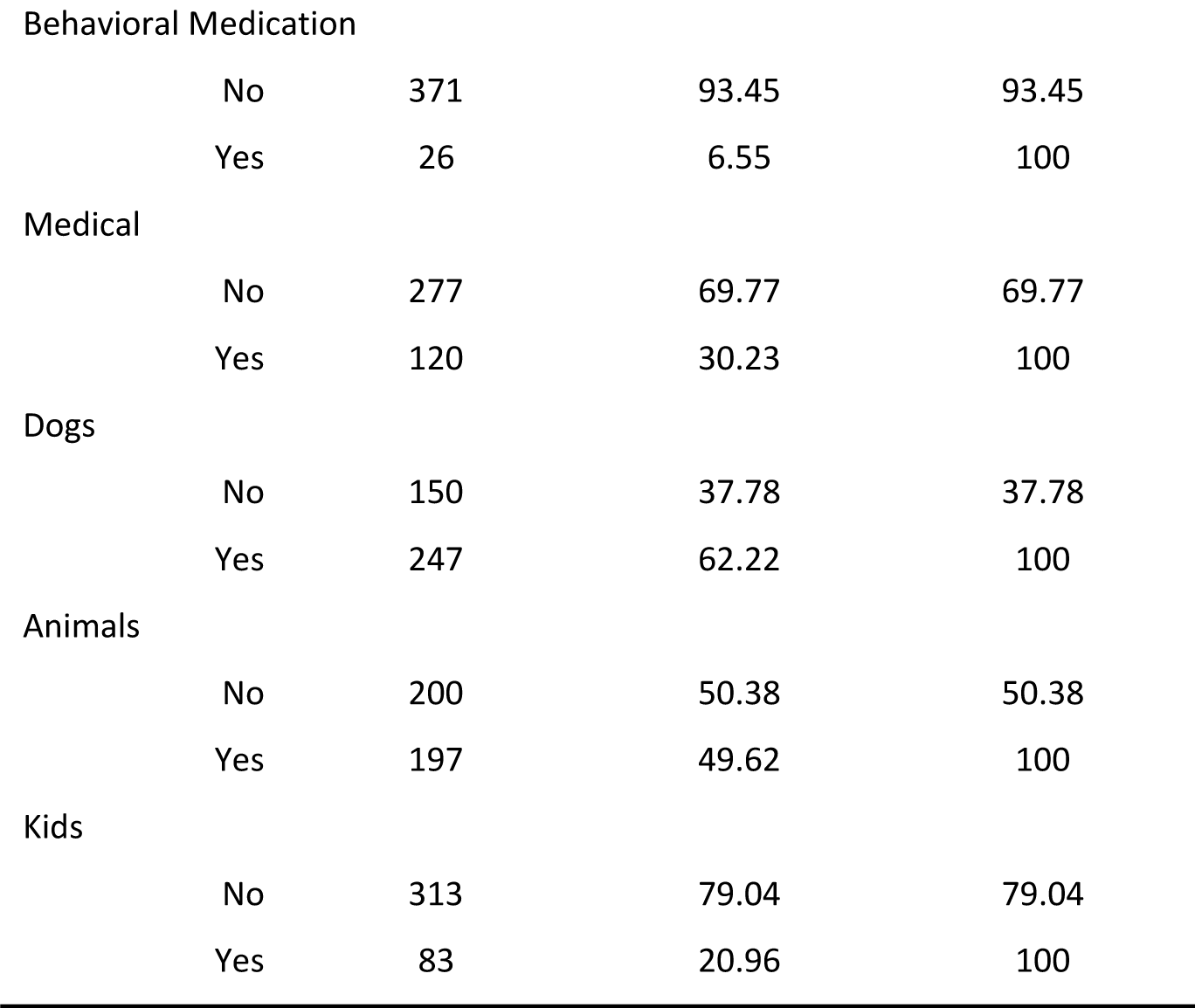
Descriptive statistics of cohort.

| Variable | Frequency | Percent | Cumulative<br>Percent |
| --- | --- | --- | --- |
| Sex |  |  |  |
| female | 189 | 47.61 | 47.61 |
| male | 208 | 52.39 | 100 |
| Neuter |  |  |  |
| Fixed | 365 | 92.17 | 92.17 |
| Intact | 31 | 7.83 | 100 |
| Work |  |  |  |
| No | 381 | 95.97 | 95.97 |
| Yes | 16 | 4.03 | 100 |
| Compete |  |  |  |
| No | 380 | 95.72 | 95.72 |
| Yes | 17 | 4.28 | 100 |
| Purebred |  |  |  |
| No | 221 | 55.81 | 55.81 |
| Yes | 175 | 44.19 | 100 |
| Pitbull |  |  |  |
| No | 338 | 85.14 | 85.14 |
| Yes | 59 | 14.86 | 100 |
| Acquire Place |  |  |  |
| Breeder | 109 | 27.53 | 27.53 |
| Other | 104 | 26.26 | 53.79 |
| PetStore | 13 | 3.28 | 57.07 |
| Rescue | 25 | 6.31 | 63.38 |
| Shelter | 145 | 36.62 | 100 |
| Other House |  |  |  |
| No | 176 | 55 | 55 |
| Yes | 144 | 45 | 100 |
| Behavioral Diagnosis |  |  |  |
| No | 275 | 69.27 | 69.27 |
| Yes | 122 | 30.73 | 100 |

|  |  |  |  |  |
| --- | --- | --- | --- | --- |
| Behavioral Medication |  |  |  |  |
|  | No | 371 | 93.45 | 93.45 |
|  | Yes | 26 | 6.55 | 100 |
| Medical |  |  |  |  |
|  | No | 277 | 69.77 | 69.77 |
|  | Yes | 120 | 30.23 | 100 |
| Dogs |  |  |  |  |
|  | No | 150 | 37.78 | 37.78 |
|  | Yes | 247 | 62.22 | 100 |
| Animals |  |  |  |  |
|  | No | 200 | 50.38 | 50.38 |
|  | Yes | 197 | 49.62 | 100 |
| Kids |  |  |  |  |
|  | No | 313 | 79.04 | 79.04 |
|  | Yes | 83 | 20.96 | 100 |

We classified dogs reported to be Pit Bull or Staffordshire Terrier/AST as Pit Bull-type dogs, which made up 15% of our cohort. The Principal Components Analyses (PCA) below and further evidence discussed in the Supplementary Text show patterns that are consistent with known Pit Bull classification rates and percent breed makeups^24^.

Owners provided dog cheek swabs for DNA isolation. We used custom TaqMan™ quantitative Polymerase Chain Reaction (qPCR) assays to genotype SNPs at 20 markers associated with problem behaviors in our mapping studies (Methods; Table 2 and Suppl. Table S3). The markers were taken from the SNP platforms used in the genome scans and are assumed to be in linkage disequilibrium (LD) with causal variants in the tagged risk haplotypes. All allele frequencies, but one, were in Hardy-Weinberg equilibrium. The Chr1A marker was detected as two states rather than three and was thus analyzed as binary. No DNA copy number variant has been described at this locus, but it remains possible the binary genotype could reflect the presence and absence of a structural variant.

**Table 2.**
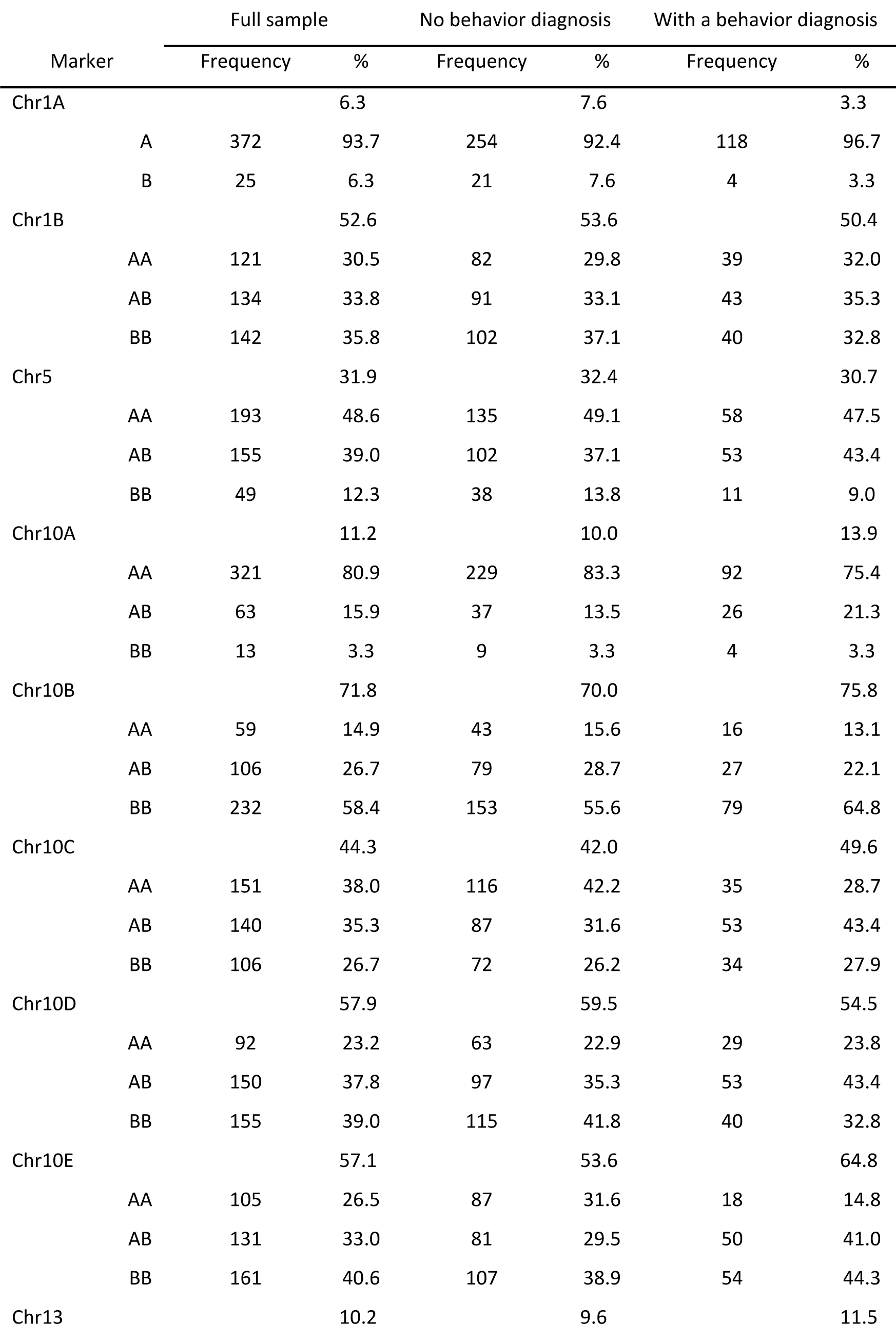
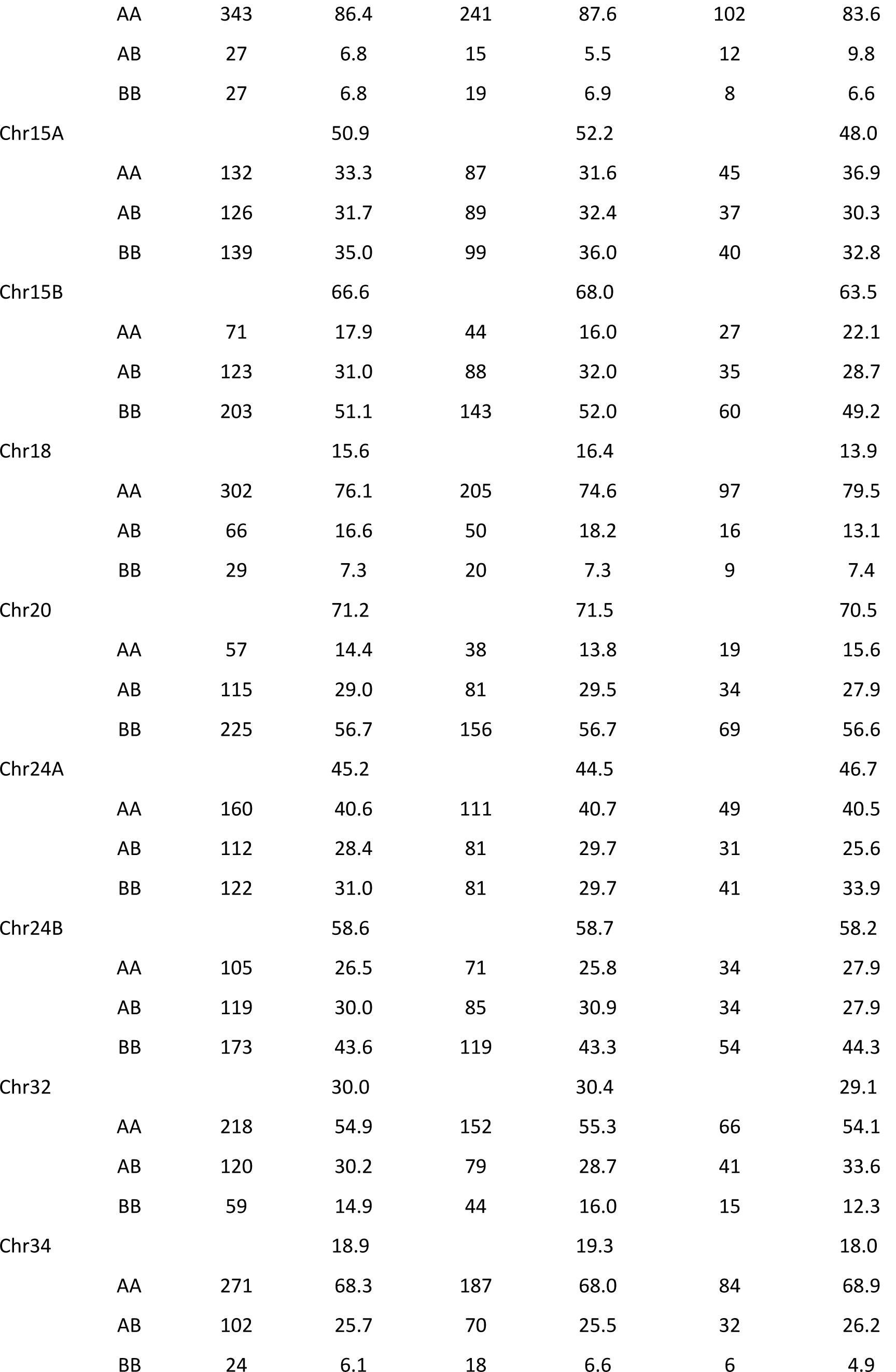
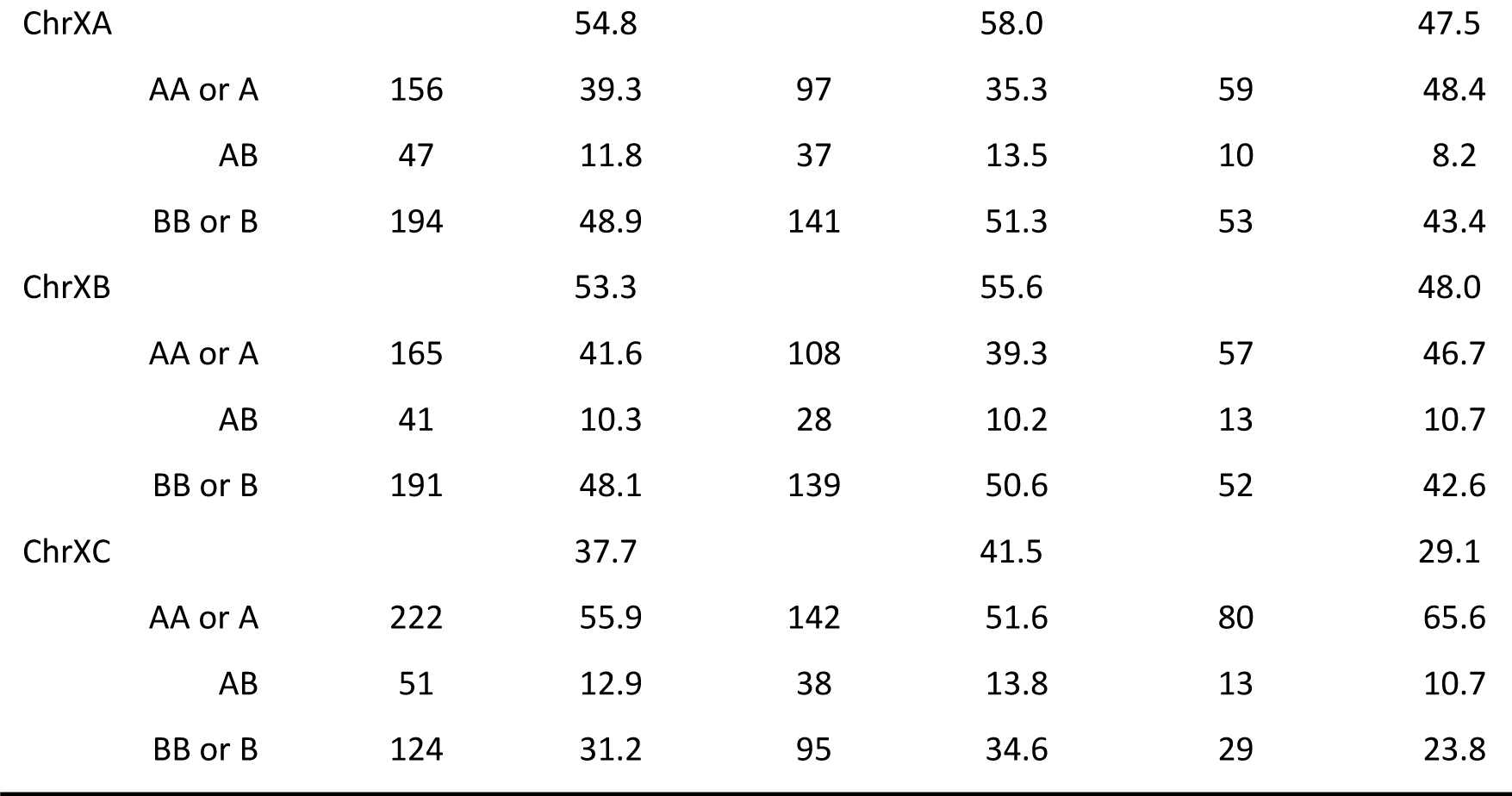
Allele frequencies for sample and diagnosis classes.

### Variable association and correlation analyses

We created association tables to evaluate the relationships of all predictor variables. Those include paper questionnaire variables, descriptive C-BARQ variables and marker genotypes. Since variables were continuous, binary or multilevel categorical, we estimated correlations using different methods (Suppl. Table S4). Figure 1 shows pairwise associations of all predictor variables (see Suppl. Fig. S1 for numerical values). Many predictable correlations are evident. Dogs with medical ailments tended to be older. Neuter status was correlated with the dog function and source. The dog source was strongly associated with other variables such as the age the dog was acquired, neuter status, whether the dog lived in another household, and Pit Bull-type status. We expected these associations due to the nature of different sources.

**Figure 1.**
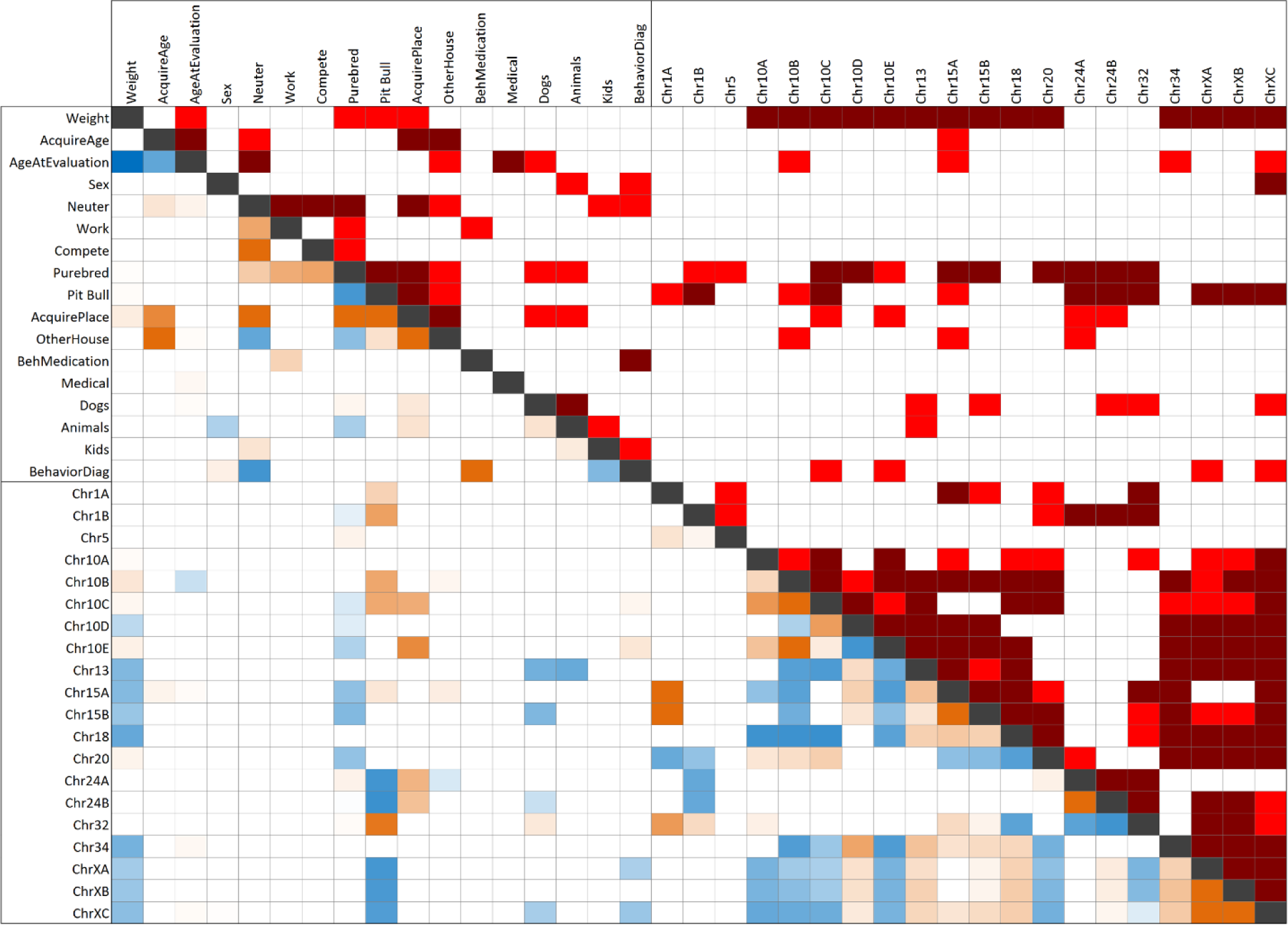
Pairwise association of questionnaire variables and genetic markers. Significance test is shown above the diagonal line and effect size and direction below (odds ratio for categorical variables and estimate ratio for continuous variables). SNP alleles are given according to the CanFam3 nomenclature: Reference allele is A and Alternative is B (A/B should be considered arbitrary assignments without regard to population frequencies or ancestral/derived status). Genetic marker significance test and correlation are for AA vs BB. In the top right, red denotes significant association, p≤0.05; and dark red is significant association, p≤0.001 (for actual p-values, see Suppl. Fig. S1). In the bottom left, red is positive association and blue negative. Values are colored in a gradient from red to blue according to their value. Only values with a significant association are displayed.

Mirroring trends of American Kennel Club breed popularities in the past two decades^19,28,29^, we observed dogs acquired from pet stores tended to be smaller in size. There was a negative association between having children in the home and having a behavioral diagnosis (discussed in Suppl. Text). Female sex was associated with any behavioral diagnosis, whereas males had increased risk for aggression directed at familiar dogs (see modeling results below). There is no consensus on the effects of sex on canine anxiety/separation traits. Females are known to have increased risk of developing fear of unfamiliar humans and dogs^20,30^. Intact females have increased fear of dogs compared to intact males, but levels are increased further – and the sexes are indistinguishable – when they are neutered^20^. Males are at increased risk of being more aggressive than females^30-35^. Here, neutering of both sexes was positively correlated with behavioral diagnosis, consistent with previous reports^20,35^. The modeling analysis has additional detail for neuter status, but we do not stress this variable because only a small percent of our cohort was intact.

We observed correlations between markers on different chromosomes, such as 10, 18, 24, 32 and X. This is presumably due to genetic stratification across breeds, but it should be noted that the present loci are likely to be enriched for admixture^26^ and possibly selection in early domestication^11,12^ (may not reflect vertical breed-relatedness). Supporting this, correlated markers were not correlated with the same traits. One consistently-detected association was between genetic markers and small body size (using weight as a proxy). We previously reported this and the association of small size and problem behaviors^11,12^, which are supported by behavioral^17-20^ and other genetic evidence^22^. Pedigree breed and Pit Bull-type represent reduced variation, and, thus predictably, showed association with subsets of markers.

To test for cohort deviations and overrepresentations of traits, we estimated C-BARQ trait associations through correlation analysis (Suppl. Fig. S2). The findings raised no such concerns about the cohort. The observed relationships across C-BARQ behavioral traits reinforce what has been described in detail^16,19,36^. For instance, this analysis and our genome scans showed a strong relationship between fear and aggression^11,12^. We interpret this to mean aggression frequently stems from fear^12^, a finding consistent with behavioral studies of canine aggression^17^.

### Principal Components Analyses

We carried out PCA of genotypes and C-BARQ response variables (Fig. 2). PCA allows visual representation of association or sampling bias: data points which cluster together are more similar than those further apart. Figure 2A shows PCA of genetic markers, for which 32.4% of the variance is explained within the first two dimensions. Some markers on the same chromosome clustered together due to LD, such as a group on chromosome 10. Others did not cluster together, such as the three at the large X chromosome region. This indicates they can be present on different haplotypes (for demonstration at the present X chromosome locus, see Figs. 6 and 8 in ref.^12^).

**Figure 2.**
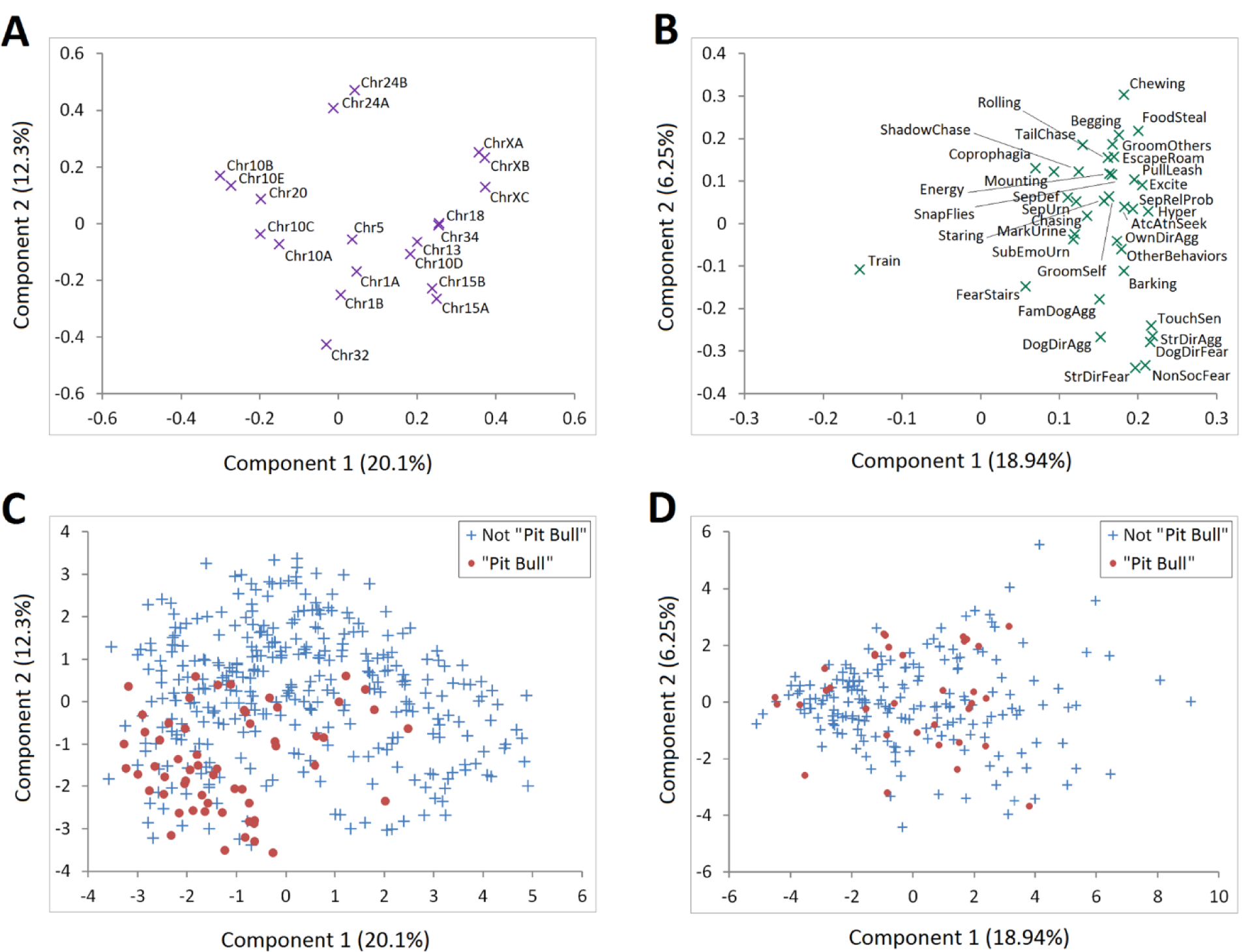
Principal components analysis (first two components). (A) genetic markers, (B) C-BARQ behavioral traits, (C, D) genetic markers and C-BARQ behavioral traits, respectively, with Pit Bull-type classification.

Figure 2B shows PCA of the C-BARQ response variables, for which 25% of the variance was explained in the first two dimensions. This plot is consistent with our previously reported correlations between fear and aggression traits^11,12^. We further evaluated the uniformity observed in our association analysis among the Pit Bull-type dogs. To accomplish this, we plotted the PCA data for genotype and C-BARQ scores, but colored according to Pit Bull status (Fig. 2C/D). A cluster of Pit Bull-type samples in the lower left side of the plot indicated those dogs are genetically more homogeneous with each other and different from the other dogs in the cohort. Examination of those dogs with pedigree dogs suggests the more homogeneous dogs are purer Pit Bulls. That is, the more tightly-clustered these dogs are, the more similar they are to breeds closely related to Pit Bulls (Suppl. Text). We similarly considered sex, neuter status, pure pedigree, mixed breed, behaviorally diagnosed, and other non-behavioral medical ailments (Suppl. Figs. S3, S4). Several of those plots showed deviations from randomness. For comparison to Pit Bull-type dogs, we generated plots for common pedigree breeds in our cohort: the combined members of the retriever group, German Shepherd Dogs and Rottweilers (Suppl. Fig. S5).

### Behavioral diagnosis prediction based on genetic markers

We next tested whether our set of 20 markers could predict the risk of dogs having a behavioral diagnosis. We used logistic regression models to evaluate our cohort of 397 total dogs, 122 of which had a behavioral diagnosis (incl. diagnosed Pit Bull-type, n=20; and medicated for behavior, n=26). The models considered all genetic predictors simultaneously, thus accounting for (but not estimating) their correlations. The results showed a set of five markers on three chromosomes (10, 13 and X) can predict a behavioral diagnosis (Fig. 3; p-values given in Suppl. Fig. S6). In most cases, the loci associated with any behavioral diagnosis were different from those associated with a specific diagnosis. This was possible because the tests of any diagnosis were done on the full cohort, but the tests for specific diagnoses were done on only the subset of dogs with any diagnosis. The top candidate genes at those loci are *MSRB3* and *HMGA2, ANGPT1* and *IGSF1*, respectively^12^. [The *RSPO2* haplotype associated with canine coat traits is near, but distinct from, the chr13/*ANGPT1* risk haplotype^11^.] There are multiple haplotypes at the second chromosome 10 risk locus (chr10B-E markers) and those are associated with various morphological and behavioral traits^7,10,12^. Subsets of these traits can be correlated in some breeds, such as small size, floppy ears and increased fear, anxiety and aggression. The chromosome 13 risk haplotype is associated with multiple behavioral traits, including increased fear, anxiety and aggression traits, as well as smaller size. Lastly, X chromosome locus markers near *IGSF1* are associated with fear, anxiety, aggression and body size traits, and markers near *HS6ST2* are associated with sociability^10-12^. Further behavioral analyses are necessary, but the first implication is that fear, anxiety and aggression are the most important emotional or personality traits associated with a clinical diagnosis of problem behaviors.

**Figure 3.**
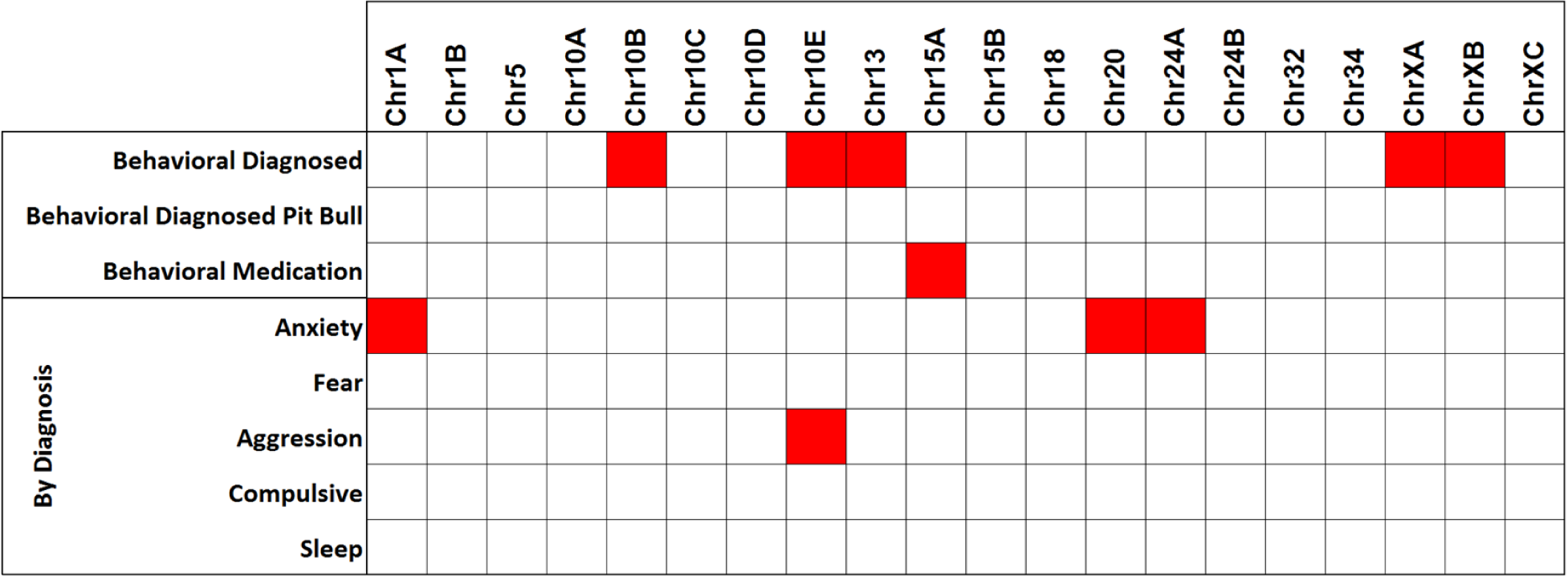
Diagnostic prediction. Top shows significant marker prediction of a behavioral diagnosis and medication usage (p-values are given in Suppl. Fig. S6). Bottom shows significant marker prediction of specific behavioral diagnoses.

Genetic markers which are strongly correlated tend to have redundant effects due to collinearity. However, none of the marker correlations present in the association tables and PCA plots reached significance. For example, the marker chrXC was not significantly associated with chrXA/B although they clustered closely in the PCA plot (Fig. 2A). No marker was associated with Pit Bull-type among behaviorally diagnosed dogs, indicating the breed is not unique or behaviorally-defined by any single variant in our panel. The allele of *IGF1* (chr15) that explains the most variance in small body size in dogs^9,37^ predicted dogs currently on medication for their behavioral diagnosis.

In terms of specific diagnoses made by a clinical behaviorist, we detected three significant marker associations with anxiety disorder: chr1 near *ESR1*, chr20 near *MITF* and chr24 near *RALY, EIF2S2* and *ASIP*. No marker was predictive of a fear diagnosis but a chr10 marker between *MSRB3* and *HMGA2* was associated with diagnosis of aggression. We did not detect associations with compulsive behavior and sleep disorder diagnoses. This was not surprising because the frequencies of those diagnoses were very low in our cohort and, therefore, only very large effects could have been detected. Our genetic testing results are consistent with reporting by the veterinary behavioral field^38,39^, and suggest clinical relevance and, presumably, broader use.

### Statistical modeling of C-BARQ behavioral traits

We used statistical modeling to determine the relevance and effect direction of each predictor variable – from the paper questionnaire and genotype markers – for each C-BARQ variable. We applied three modes. The full model mode (FMM) included all predictive variables together. FMM offers risk estimation incorporating covariation introduced by other variables in the model. The individual model mode (IMM) included each predictive variable individually. The IMM does not take any covariance into account and offers risk estimation independent of other predictors. The fixed threshold case-control mode (FTCCM) stratified risk according to trait severity, but has increased uncertainty. As severity increases, there are fewer cases and power decreases. Therefore, each mode has its own inferential application that requires further evaluations of utility. Due to the statistical power that can be achieved in our sample, it was not feasible to include individual breeds in our models. We believe this is unnecessary because our risk alleles were mapped by interbreed GWA in multiple cohorts and are thus common across diverse breeds rather than representing specific breeds or breed groups^26^.

Overall, the IMM detected more significant associations than the FMM (Figs. 4/5; p-values given in Suppl. Figs. S7/8). The most consistent predictor variables were: i) having a behavioral diagnosis (16/36 FMM vs. 15/36 IMM), which consistently increases risk of problematic behavior for several traits; ii) participating in competitive sports (10/36 FMM vs. 9/36 IMM), which reduces risk for most problematic traits except for familiar dog aggression; iii) age of acquisition (9/36 FMM vs. 6/36 IMM); and iv) age at evaluation (9/36 FMM vs. 11/36 IMM). The latter two are associated with both reduced and increased risks of different traits. Consistent with previous behavioral studies^12,19^, we found that larger dogs are considered to be more trainable. Working dogs had higher risk of separation-related problems, increased energy and coprophagia (FMM only). The energy trait, and possibly separation, is consistent with a previous study of Swedish military working dog temperament using C-BARQ phenotypes^40^. SMWDs which passed temperament tests were more hyperactive/restless – which we showed here to be correlated with energy (p<0.001, Fig. S2) – than those who did not, and were, on average, left home alone more hours per day in their first year of life. Dogs with non-behavioral medical ailments had a reduced risk of displaying many problematic behaviors. They also showed an increased risk of aggression directed at familiar dogs (FMM and IMM) and coprophagia (IMM only).

**Figure 4.**
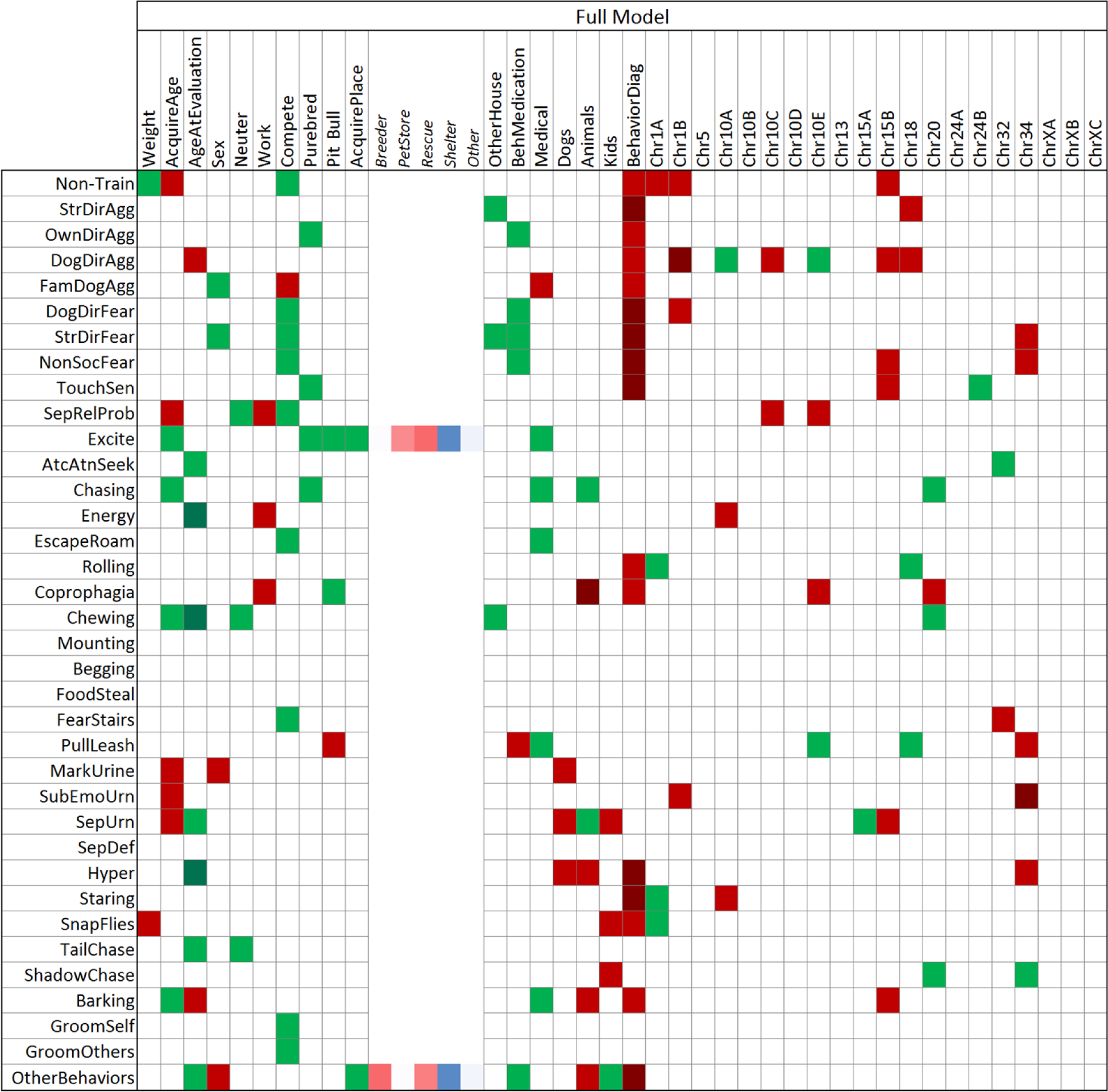
Full Model Mode (FMM). Generalized linear model associations of C-BARQ behavioral traits by questionnaire and genetic markers were evaluated together. Each behavioral trait was modeled but only significant effects are highlighted. SNP alleles are given according to the CanFam3 nomenclature: Reference allele is A and Alternative is B (A/B should be considered arbitrary assignments without regard to population frequencies or ancestral/derived status). Green denotes decreased risk and red increased risk of the A vs. the B allele. A darker shade of green or red denotes significant at a Bonferroni level adjusted by trait. Actual p-values are given in Supplementary Figure S7. When the effect of place acquired (AcquirePlace) is significant, the Least Square Mean estimate of each of its levels is shown in the columns to its right; color gradient is arranged from lowest to largest.

**Figure 5.**
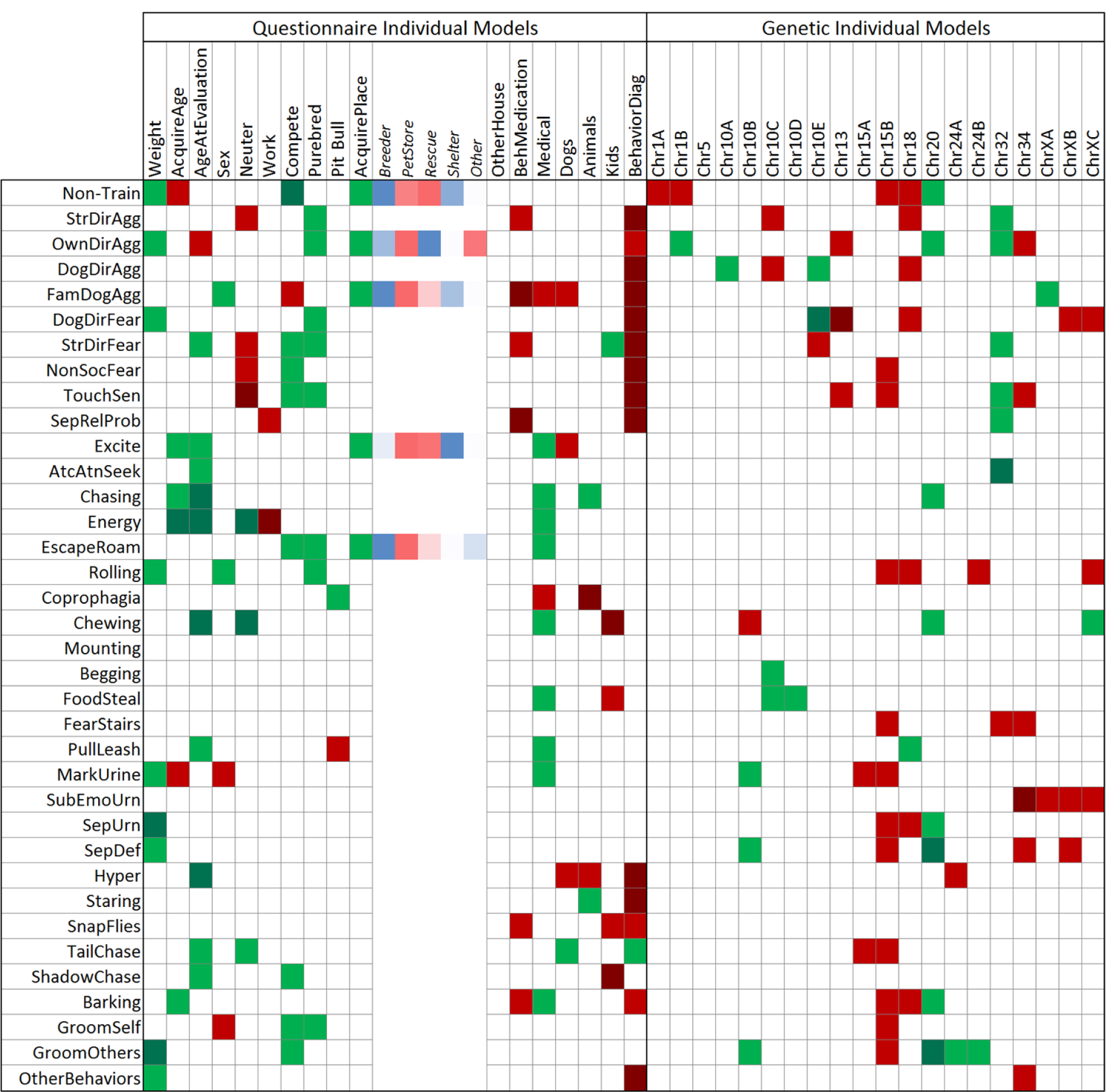
Individual Model Mode (IMM). Generalized linear model associations of C-BARQ behavioral traits by questionnaire and genetic markers were evaluated individually. Each behavioral trait was modeled but only significant effects are highlighted. SNP alleles are given according to the CanFam3 nomenclature: Reference allele is A and Alternative is B (A/B should be considered arbitrary assignments without regard to population frequencies or ancestral/derived status). Green denotes decreased risk and red increased risk of the A vs. the B allele. Green denotes decreased risk and red increased risk. A darker shade of green or red denotes significant at a Bonferroni level adjusted by trait. Actual p-values are given in Supplementary Figure S8. When the effect of acquired place (AcquirePlace) is significant, the Least Square Mean estimate of each of its levels is shown in the columns to its right; color gradient is arranged from lowest to largest.

Having other non-canine animals in the home was associated with reduced risk of chasing and increased risk of coprophagia and hyperactivity (FMM and IMM). Having other dogs vs. other animals in the home only overlapped for increased hyperactivity (both modes). Dog body size had two behavioral trait associations in the FMM and nine in the IMM. This hints that small body size is a relatively good predictor when more information is unavailable, consistent with previous reports^17-19^. Pit Bull-type designation was not predictive of aggressive behavior, but reduced risk of coprophagia and excitability (FMM), and increased risk of leash pulling (both modes). Having children in the home increased the risk of snapping at flies and shadow chasing, and reduced the risk of stranger-directed fear (IMM) and other stereotypic behavior (“Other Behaviors”; bizarre, strange, or repetitive behavior). The source of acquisition from was predictive of excitability, some types of aggressive behavior, and trainability (IMM). Dogs obtained from a shelter or from a breeder tended to have lower risk of problem behaviors than dogs from pet stores. That is consistent with the fact that dogs purchased at pet stores have increased risk of problem behaviors compared to dogs from non-commercial breeders^41^. This could be confounded by small body size, which is genetically associated with problem behaviors^11,12^. The last two decades have experienced a trend of increased popularity of smaller pedigree breeds^19,28,29^.

The most consistent genetic effects came from two body size loci^9^: the chr15B marker at the *IGF1* locus (6/36 FMM vs. 12/36 IMM) which always increased risk, and chr34 near *IGF2BP2* (6/36 vs. 6/36), which, in the majority of cases, increased risk of multiple problem behaviors. The most consistent genetic effect predicting fear and aggression was chr18 at the *GNAT3/CD36* locus, as previously reported^11,12^. In both FMM and IMM, the chr20 marker near *MITF* (4/36 FMM vs 8/36 IMM) predicted reduced risks of inappropriate chewing and chasing. In both modes, chr10E consistently predicted reduced risk of aggression directed at unfamiliar dogs, but mixed effects for other traits. Some markers, like chr32 near *RASGEF1B*, showed an effect across several traits but only in a single mode. Chr10B, chr10D, chr13, chr24A and all three chrX loci were only significant in IMMs. The C-BARQ trait with the most genetic predictors was aggression directed at unfamiliar dogs (6/20 FMM vs. 4/20 IMM). Interestingly, owner-directed aggression had 5/10 genetic predictors in IMM but none in FMM. Both modes consistently showed chr10A near *LRIG3* predicted aggression directed at unfamiliar dogs. In summary, genetic testing consistently predicted multiple C-BARQ traits, including in the areas of fear, anxiety and aggression.

We used the FTCCM to test the robustness of significant predictors and determine how association patterns are affected as behavioral C-BARQ traits are stratified by severity. We set fixed threshold values for each C-BARQ trait at the 50^th^, 75^th^, 90^th^ and 95^th^ quantile levels and deemed those above the threshold as cases. We analyzed the data as a binary response variable. We created models using logistic regression with a stepwise forward selection. Overall, FTCCM models showed a similar pattern as the FMM and IMM generalized linear models (Suppl. Figs. S9-12). As fixed thresholds were raised with concomitant loss of power, we detected some interesting effects. Weight, age and age at evaluation decreased their relevance as the threshold increased. As was observed in FMMs and IMMs, having a behavioral diagnosis consistently predicted problematic behavior and dogs with non-behavioral diagnoses had lower risk of some problem behaviors.

As expected, FTCCM models were the least sensitive. Genetic marker performance for FTCCM models was not as robust as in FMMs and IMMs. The exception was chr5 (near *SHISA6*), which exhibited the least variability in the PCA (Fig. 2A) and was not significant in the FMM and IMM. Chr5 was mapped for escaping and chasing^11^, and here was associated with escaping in the FTCCM. ChrXB predicted milder cases of urine marking. Chr10E, chr18, and chrXC were the most relevant for fear and aggression traits. Chr10E was most relevant when the threshold was lower (50 and 75^th^ quantiles), suggesting it segregated milder cases of dog directed fear. Chr18 and chrXC were most relevant for detecting intermediate cases when thresholds were set to 75-90^th^ quantiles (aggression directed at unfamiliar humans and fear directed at unfamiliar dogs, respectively). Three markers had increased relevance for detecting problematic behavior of greater severity, 90-95^th^ thresholds: chr1A and chr34 for touch sensitivity, and chr20 for separation-related defecation. Chr32 was associated with increased trainability (90-95^th^ thresholds). Curiously, the chr32 trainability association was not present in the FMM and IMM, but reduced fear of stairs was observed in all three models and at all FTCCM thresholds. Pit Bull-type dogs were not associated with any fear or aggression trait in the FMM or IMM models. In the FTCCM, they showed reduced risk of owner-directed aggression only at the 75^th^ quantile of severity and increased fear of unfamiliar dogs only at the 95^th^ (discussed in Suppl. Text).

## Discussion

One strength of genome scanning of breed averages is the ability to map alleles that are fixed in individual breeds^7,9,10,12^. This can complicate interpretation and validation in those breeds, but that can be addressed in other breeds and in mixed breed dogs. A drawback of the approach is that it cannot detect variants that are rare across breeds^42,43^. A second strength of interbreed mapping is that causal variants can be assumed to lie within the minimal overlap region across breeds carrying the risk haplotype^7,9,10,12^. Because meiotic recombination events happen independently in each breed, LD breaks down on both sides of causal variants and the markers tagging them. As a result, the peak intervals in interbreed GWASs tend to be much smaller than in single breed GWASs. Additional virtual fine-mapping is possible by breed-specific phasing of GWA haplotypes to further refine the minimally overlapping region^11,12^. Notably, our original GWASs were made possible by using crowdsourced C-BARQ phenotypes and unrelated genotype datasets of only partially-overlapping dog breeds. Here we tested 20 SNP markers at 13 of those loci for behavioral associations in a 397-dog cohort designed to randomly sample the community and behavioral clinic. In contrast to the GWASs, the present study had individual-level C-BARQ phenotypes and genotypes. Because of the high complexity of this work (incl. 17 behavioral traits and use of half mixed-breed dogs and half dogs representing 77 breeds) and the low power of the cohort, we consider the GWASs a first phase of discovery and this study a second phase. Our findings supported behavioral associations for all loci tested, but confirmatory studies will require a narrower scope or much greater power.

Control of population structure is critical to genetic studies of domesticated species^4,21,44^. We previously mitigated the effects of population structure by using linear mixed models and multiple cohorts with partially overlapping breed makeup in the discovery GWASs^11,12^. We also provided evidence for a subset of markers through predictive modeling in a third group of dogs with no breeds overlapping the GWASs^12^. Using many breeds and multiple cohorts with different breed makeups reduces the risk of false positives due to population structure and latent variables such as cryptic relatedness and batch effects. Here, we observed correlations between unlinked markers in our cohort. This is consistent with stratification of genetic variation across breeds. Despite the large number of breeds included and the high proportion of mixed breed dogs, this is not surprising. Breed popularity is so unbalanced that the 10 most popular breeds accounted for 50% of all 2008 AKC registrations. We cannot rule out the effects of population structure on our studies or that some behavioral variants are part of, or inextricable from, population structure^4,21,44^. However, population structure is not a critical problem here because markers that are correlated genetically are not correlated with the same traits. For instance, chr18 and chr34 correlated with several markers associated with having a behavioral diagnosis; however, chr18 and chr34 did not correlate with behavioral diagnoses in our association, prediction and modeling analyses (Figs. 1, 3-5). The same is true for chr32 and fear of stairs (see below and Suppl. Text). We considered our results in the context of high-powered clustering of breeds according to C-BARQ behavior, which found clusters are most strongly associated with body size, followed by breed relatednes^29^. PCA classifying our data according to those clusters showed partial segregation of our genetic markers but not of behavior (Suppl. Text and Figs. S13/14). Lastly, strong biological relevance of candidate genes further supports our behavioral associations^11,12^. For example, the implicated genes at the two loci most-associated with fear and aggression directed at unfamiliar humans and dogs – chr18 and chrX here – are supported by evidence in rodents of related behaviors and gene expression in the amygdala to hypothalamic-pituitary-adrenal axis^12^.

This study provided further support for our genome scans of canine behaviors^11,12^ and suggested their clinical relevance. Ten markers at eight loci were associated with having a clinical behavioral diagnosis, and a set of five of those successfully predicted a diagnosis. Among the broad corroborating evidence of C-BARQ associations (Suppl. Text), we found evidence that supports all four of the original loci replicated in a second cohort for nine fear and aggression traits (chr10, chr15, chr18 and chrX)^12^. The chr18 and chrX associations in this cohort support our original interpretation that variants at those loci are associated with fear and aggression directed at unfamiliar humans and dogs, but not with owner-directed aggression^12^. Most of the traits were further supported by the same trait or a related one^11,12^. We also provided further evidence for the GWA findings for chromosome markers 1B, 10A (very distant from 10B-E), 20, 24 and 34. While our findings lend support to many of the mapping results, most variations also had trait associations that differed from the GWASs^11,12^. This is unsurprising given the differences in design and power, and the high levels of pleiotropy known for human brain traits^45,46^ (discussed below). For example, chr32 was associated with aggression in the GWASs and here. However, that chr32 haplotype differed for several anxiety traits across the GWASs and here. More studies are necessary to determine if our GWA of breed averages or the present study with individual-level data predicted associations more reliably. We expected this work to yield more accurate data, but it was recently shown that GWA of dog body size was dramatically more powerful using breed averages than individual measures^47^.

Trait correlations should be considered carefully as they could vary across breeds or be due to environmental effects. A previous study supports the general negative correlations of trainability with energy and snapping at flies we observed^15^. However, there is evidence the trainability-energy relationship is not fixed. For example, in comparisons of breed groups, sighthounds rank at the top for both trainability and energy^30^. Trainability and energy can also be positively correlated in working dogs, but interpretation is complicated due to the effects of selection of dogs for training and of the training itself^48,49^. Among the suggestions of gene-environment interactions, we found strong correlations of behavioral phenotypes with presence of children in the home. Lastly, we note that both barking and coprophagia are more prevalent in domesticated dogs than wolves^50^. But, whereas barking seems to be an early target of human selection, the reason for coprophagia is unknown. Canine coprophagia generally involves non-autologous fresh stools^51^, which we believe is suggestive of microbiome inoculation. In humans and mice, transplantation of fecal microbiota has therapeutic effects on anxiety, depression and inflammation^52,53^. Here we found coprophagia was associated with both behavioral and non-behavioral medical diagnoses. Further studies are necessary to determine if coprophagia is simply correlated with illness or if it could be an adaptation with therapeutic benefits.

Our prior^12^ and present findings in dogs are suggestive of pleiotropy (Suppl. Text and Table S15). That is also strongly supported by comparative genetic analyses of dog behavioral GWASs^11,22^ and consistent with the rapidly growing evidence of widespread pleiotropy of behavior in humans^45,46^. For instance, we showed risks of many dog problem behaviors are associated with specific genetic variants known to cause small body size (*IGF1, IGF1R, IGF2BP2* and *HMGA2*) and that protection against problem behaviors is conferred by the large-size *IGSF1* haplotype^12^. In the present study, we tested all but the *IGFR1* locus and provided further support for these relationships. Several canine behavioral traits associated with reduced body size are correlated with each other^18^, and those effects were consistent with our genome scanning results^12^. Veterinary behaviorists have previously shown that small dog size is associated with problem behaviors^18,20,29^. A study of German Shepherds showed that drug-detection training results in immediately increased levels of circulating IGF1; and this effect is potentiated in dogs that have undergone six months of training vs. none^54^. While German Shepherds are fixed for the non-small body size allele, this finding suggests physiological relevance for trainability – which is one of the traits we showed to be negatively associated with the small body size allele of *IGF1*.

In humans, there are many genetic correlations between height and psychiatric, behavioral and personality traits, including neuroticism^55^, risk tolerance^56^ and smoking cessation^57^. There is also strong evidence that body size is associated with differences in brain structure in humans and dogs, and that those have functional effects (most commonly reported in the area of cognition)^11,58^. Both Insulin/IGF signaling and downstream pathways (here incl. *IGF1, IGF2BP2* and *IGSF1*, and *HMGA2*, respectively) have important roles in brain development^59,60^. Presumably correlations of dog body size and behaviors also involve physiology and psychology^12,59^. Our interpretation is that the behavioral genetic pathways we mapped are conserved at least across mammals. However, although body size has been under selection in both humans and dogs, the biology and genetic architecture are dramatically different^11,61,62^. Dog body size is mostly explained by a handful of variations of moderate-to-large effect sizes, whereas humans have countless variations with weak effects. It seems likely this would be reflected in the pleiotropy of those variations.

Our findings for Pit Bull-type dogs have three uncertainties (Suppl. Text). First, the designation of Pit Bull-type dogs is based on visual appearance and the expectation that mean AST content was ∼40-50%^24^. If that AST makeup were correct, we believe our study of these dogs is justified since the correct classification rate is 76% for dogs as low as 25% AST^24^. Pit Bull-type dogs have increased genetic diversity because they represent multiple breeds and because they are commonly mixed with other breeds. As a result, true Pit Bull effects are distorted and diluted, and the power to detect them is reduced. Secondly, our interbreed behavioral GWASs could not have identified risk variations that are common in Pit Bull-type dogs but otherwise rare. And thirdly, our Pit Bull-type sample is probably under-represented for dogs know to be exceptionally aggressive or to be very successful in dog fighting (which would be associated with increased risk of dog attacks and criminal behavior^63^). Both modes of the generalized linear modeling showed only a single trait association: increased leash pulling. The FTCCM mode detected decreased owner-directed aggression at the 75^th^ quantile of severity and increased unfamiliar dog-directed fear only at the 95^th^. A previous study of C-BARQ aggression traits in approximately 3,800 dogs included Pit Bull-type dogs as defined here^17^. It showed they have reduced risk of owner directed aggression, as we observed, and increased risk of aggression directed at dogs – but not humans. It is unknown whether the latter 11.5% of Pit Bull-type dogs with increased dog-directed aggression also had increased fear of dogs. If that were the case, it would explain our observation of extreme dog-directed fear in a small subset of this breed type. However, our community sample of Pit Bull-type dogs showed they are not more aggressive or more likely to have a behavioral diagnosis than other dogs. This does not support reliance on breed-specific legislation to reduce dog bites to humans^63^. As our genetic findings were restricted to known aggression variations that have large effect sizes across breeds, it is necessary to identify and understand the effects of rarer loci that increase risk of dangerous behavior.

Population structure is the most challenging aspect of genetics in domesticated species. This can be addressed by the design of future confirmatory studies in dogs. Those will also make it possible to measure the proportion of the trait variance explained by single and combinations of variations^13^. We successfully applied such concepts to canine osteosarcoma, including using the Intersection Union Test to perform a type of meta-analysis, modeling polygenic risk within and across breeds, and validating one breed model in a separate sample^6^. It is currently not feasible to conduct well-powered mapping in the several hundreds of existing dog breeds. However, it is possible to study the most popular breeds (in the US, 10 breeds account for 50% of all AKC registrations). Alternatively, behavioral genome scans could be performed in phenotyped mixed-breed dogs^64^ and those haplotypes could be characterized in those and pedigree dogs. Other factors that are difficult to consider in this work and should be addressed in follow-on studies are dog sampling bias, and effects of owner personality and socioeconomics. Here we showed our cohort is representative of the community. However, we assume that owners of dogs with behavioral diagnoses have a higher socioeconomic status than average because they were recruited through an academic veterinary hospital. Many canine behavioral variants may require environmental stimuli for a behavioral phenotype to manifest. Owner personality does not necessarily increase the risk of owner-directed aggression^65^, but owner personality and psychiatric traits are correlated with increased rates of fear, anxiety, aggression and other traits^66,67^. Caution must be used in interpreting the association of small body size with problem behaviors. Small dogs, as a group, may have different owner and other environmental characteristics compared to larger dogs (e.g., physical and social characteristics of home and neighborhood, amount of time spent alone, and levels of physical and mental exercise). Especially when experienced early in life, stress is associated with increased risk of mental health disorders in humans and dogs^68^.

## Conclusions

This work provides further support for our interbreed genome scans of dog behaviors, and expands the relevance to mix-breed dogs. In addition to its utility to address unmet veterinary needs, there is a strong case for using dog models to understand human psychiatric disorders^12,19,69-72^. As we previously reported^12^, small body size was associated with many problem behaviors. The results support our previous findings that fear and aggression traits directed at dogs and unfamiliar humans cluster together and with non-social fear^12^. We previously noted that owner-directed aggression lies outside the latter cluster of traits and here found evidence suggesting it may be more closely associated with anxiety traits rather than fear. An important finding was Pit Bull-type dogs in our community sample, as a group, were not more aggressive or likely to have a behavioral diagnosis than other dogs. As the nascent field of canine behavior advances, it will be important to better account for human influences on dog behavior. Our results showed genetic screening of canine behavior is feasible and suggest it may be useful for owners, breeders, shelters, working dog institutions and veterinarians. However, we advise caution with direct-to-consumer tests until there is a better understanding of the behavioral risks associated with these alleles and others which may be common in few breeds, but undetectable in our interbreed approach.

## Materials and Methods

### Subject recruitment and questionnaires

Dog owners residing anywhere in the US were recruited to participate through public announcements. One was targeted to behaviorally diagnosed dog patients at the Behavioral Clinic in the Veterinary Medical Center at The Ohio State University (OSU). Due to regulatory restrictions, their medical records were not used here. Internal announcements to general staff and students were made at Nationwide Children’s Hospital, the Animal Sciences Department at OSU and the Blue Buffalo Clinical Trial Office, Veterinary Medical Center at OSU. Participants were encouraged to invite other dog owners and to submit samples from multiple dogs in their household. We excluded from our study dogs younger than 4 months old or living with the current owner for less than one month. Directly-related dogs (siblings, parents) were excluded unless the owners indicated they had very different behavior profiles (e.g., if one sibling was behaviorally diagnosed but another had no problem behaviors). We excluded dogs suggestive of aggression during cheek swabbing (which accounted for a total of one excluded dog). After a prescreening, a kit was mailed to the address provided by the participant. This kit included a DNA collection kit (see below), a paper questionnaire to be filled by the owner about their dog, instructions on how to fill the C-BARQ online questionnaire, a study consent form to be signed by the owner, and shipping materials and prepaid envelope for sending the sample to us. Owners were instructed to complete the C-BARQ online questionnaire developed and managed at the University of Pennsylvania by J.A.S. Only dogs recruited for this study were used from the C-BARQ data. In addition, a paper questionnaire (available as Suppl. Data 1) was included to capture additional details (e.g., limited household information, and behavioral and medical conditions of dogs). Subjects with missing information were excluded. Complete participation was compensated with a $5 gift card.

All dog samples and information were acquired under an approved IACUC protocol from OSU (Protocol number 2017A00000116). All methods were performed in accordance with the relevant guidelines and regulations. Owner questionnaires were reviewed by the OSU IRB board and declared exempt. All regulatory requirements of the study were approved by the BBCTO at The OSU College of Veterinary Medicine. All laboratory work was performed at The Research Institute at Nationwide Children’s Hospital, which reviewed the proposed study and determined it to be IACUC and IRB exempt.

### DNA isolation and genotyping

DNA samples were collected using one Performagene cheek swab (DNAGenotek Inc. Canada). Samples were incubated for 4-12 hours at 50°C for nuclease deactivation, stored at room temperature and processed in batches following the Performagene PG-AC1 protocol. DNA concentrations were determined using a NanoDrop spectrophotometer (Thermo Fisher Scientific).

We previously reported canine interbreed behavioral GWASs^12^. That was achieved using C-BARQ breed stereotypes of behavior and two genome wide SNP genotype datasets^7,10^. Those studies were expanded to include other C-BARQ traits and to add a third SNP genotype dataset^8^ (manuscript in preparation). For the present work, we selected 20 of those SNP markers for follow-up and modeling (Table 2; Suppl. Table S2). The loci were primarily selected for veterinary clinical relevance and prioritized by GWA detection in multiple cohorts (8 were present in 3 cohorts, 8 in 2 and 4 in 1). The latter four were selected for the biochemical or biological relevance of candidate genes. Some loci have single markers and others multiple. Most of the latter are commonly in LD across breeds, but GWA risk alleles at the second chr10 locus (B-E) and the X locus can be present on the same or different haplotypes depending on the breed (discussed in^12^; ^7,10^). Because the three GWA cohorts were not genotyped on the same SNP platform, we selected the present markers from the dataset with the highest resolution at each locus. These markers were genotyped using custom TaqMan™ qPCR genotyping assays manufactured by Applied Biosystems (Thermo Fisher Scientific). Probes were designed using their proprietary probe design tool using sequences from the CanFam3.1 UCSC Genome browser and considering any other adjacent SNPs included at the CanFam3.1 assembly included in the Broad Improved Canine Annotation v.1 ^73^. TaqPath ProAmp Master mix was used. Assay conditions we optimized and qPCR assays were run on 96 well plates on an Applied Biosystems 7500 Real Time PCR instrument using the standard protocol. Genotype data are available as Supplementary Data 1.

### Statistical Analysis

#### Descriptive statistics for correlation studies and PCA analysis

All statistical analyses in this work are reported using the CanFam3 nomenclature for SNP alleles: Reference is A and Alternative is B. The analyses were performed on SAS Enterprise Guide v.7.1 (SAS Institute, Cary, NC) running base SAS v.9.4 and SAS/STAT v.14.1. Variables included in this study were of three types: continuous variables (Suppl. Table S4), binary variables and multilevel categorical variables. Each of those types of variable has its inherent properties which were evaluated and analyzed on a case by case basis. No data transformations were necessary or implemented. Descriptive statistics were calculated using PROC MEANS for continuous variables and PROC FREQ for binary and categorical variables and PROC MIXED for combinations of binary/categorical and continuous variables. Correlations were calculated using PROC CORR for continuous variables, and PROC FREQ for binary and categorical variables. PCA was performed on the genetic markers by assuming a linear dosage effect of the alternate allele and to C-BARQ traits by assuming a linear dose response of the alternate B allele. All PCA were performed using PROC PRINCOMP. Observations with missing values were omitted in the PCA (but not in the modeling).

#### Association and statistical modeling

Association models for behavior, medication and type of behavioral diagnosis were performed using PROC LOGISTIC using a full model which included all genetic markers entered as categorical variables. Behavior, medication type and behavioral diagnosis modeling were performed only in the subset of subjects that had a formal diagnosis and those that were medicated within that subset.

Association models for C-BARQ traits and all questionnaire and genetic markers were estimated using PROC MIXED in two modes using all subjects. One mode included all predictors as a full model mode (FMM) and a second mode evaluating each predictor as an individual model mode (IMM). We estimated the Least Square Means for the “AcquirePlace” multilevel categorical variable only when it was detected as significant. We used Least Square Mean differences to determine effect directions. Effect directions were reversed for the Trainability C-BARQ trait because it is the only variable that captures a positive trait. To perform the fixed threshold case/control modeling mode (FTCCM), we used quantile values estimated by PROC MEANS for each C-BARQ trait at 50, 75, 90 and 95 percentiles to define case control status of our cohort (Suppl. Data 2). The closest score value to the quantile value above was used as a threshold and al observation with a value equal or above the threshold were designated as cases. Stepwise forward selection models were built by PROC LOGISTIC using a 0.1 threshold to determine predictors entering and staying in the model. Effects were determined by the direction of the odds ratio estimates taking the event “No”, “Intact”, “Female” and the genotype “AA” as the baseline. We considered the study exploratory and used familywise multiple testing correction^74,75^. The models FMM, IMM and FTCCM, which used the same variables, each had a different null hypothesis and family of tests. Multiple testing correction was thus based on the number of parameters per trait in each model. This p-value threshold corresponds to correction for 40 tests per trait or p ≤ 0.00125.

## Acknowledgements

We are grateful for the kind participation of owners and the contributions made by their pets. We thank the Blue Buffalo Clinical Trial Office at The Ohio State College of Veterinary Medicine for their help towards recruiting behaviorally diagnosed participants. We thank the Department of Animal Sciences at Ohio State for their help with recruiting participants from the general community. We thank the Stanton Foundation for making this work possible through their financial support.

## Author contributions

IZ and CEA designed the studies. IZ designed the statistical analyses. IZ conducted the non-clinical recruitment. MEH and MLL made the veterinary behavioral diagnoses, and, with IZ, conducted the clinical recruitment. JAS processed the C-BARQ questionnaire data and guided its interpretation. IZ performed the experimental work. IZ and CEA conducted the analyses and primary interpretations. CEA and IZ wrote the manuscript. All authors were involved in the final interpretation of the results and contributed to writing the manuscript.

## Data availability

Data are provided as Supplementary Data.

## Ethical approval and informed consent

All dog samples and information were acquired under an approved IACUC protocol from OSU (Protocol number 2017A00000116). Owner questionnaires were reviewed by the OSU IRB board and declared exempt. All regulatory requirements of the study were approved by the BBCTO at The OSU College of Veterinary Medicine. All laboratory work was performed at The Research Institute at Nationwide Children’s Hospital, which reviewed the proposed study and determined it to be IACUC and IRB exempt.

## Third party rights

None

