## Supplementary information (text, tables and figures) for "Genetic testing of dogs predicts problem behaviors in clinical and nonclinical samples"

### Supplementary information index

Supplementary Data 1 (MS Excel), paper questionnaire, C-BARQ phenotype and genotype data

Supplementary Data 2 (MS Excel), cumulative frequencies used to define Fixed Thresholds for FTCCM

Supplementary Text, Tables and Figures (this PDF)

- Supplementary Text, extended introduction and discussion
- Table S1. Cohort breed frequencies
- Table S2. Comparison of cohort breeds to US and US cities popularities
- Table S3. Study markers list with genome scan traits
- Table S4. Descriptive statistics for continuous variables
- Figure S1. Duplicate of Figure 1, but with numerical values (incl. p-values)
- Figure S2. Correlation table of C-BARQ variables
- Figure S3. PCA of C-BARQ traits isolating the following: sex, neuter status, pedigree vs. mixed breed, Pit Bull-type, behavioral diagnosis and non-behavioral ailments
- Figure S4. PCA of genetic markers isolating the following: sex, neuter status, pedigree vs. mixed breed, Pit Bull-type, behavioral diagnosis and non-behavioral ailments
- Figure S5. PCA of genetic markers separately isolating all retrievers and German Shepherd Dogs
- Figure S6. Duplicate of Figure 3, but with p-values
- Figure S7. Duplicate of Figure 4, but with p-values
- Figure S8. Duplicate of Figure 5, but with p-values
- Figures S9-12. Logistic regression with stepwise selection modeling with cases classified by trait severity at $50^{th}$, $75^{th}$, $90^{th}$ and $95^{th}$ percentile, respectively
- Figures S13/14. PCA of genetic markers and C-BARQ behavior isolating Wilson et. al 2018 clustering of breeds by C-BARQ behavior

### Supplementary Text

### Expanded introduction

#### Pit Bull-type dog behavior

The term Pit Bull does not signify a breed, but rather a group of related breeds[1-5]. Some of those are registered breeds of American Kennel Club (AKC): American Staffordshire Terrier (AST) and Staffordshire Bull Terrier. Other Pit Bull breeds are the American Bully and the American Pit Bull Terrier, which are registered by their own organizations and recognized by the United Kennel Club in the US. However, the vast majority of Pit Bull-type dogs are not registered pedigree dogs and they are most commonly classified based only on appearance[1-4]. Although the relevant AKC breeds and the UK Staffordshire Bull Terrier type dogs are no longer bred for aggression, the much more numerous American Pit Bull Terrier can be bred for fighting ("game bloodline") or not. A recent genetic study sampled 919 dogs from two dog shelters in the US states of AZ and CA[4]. 238 dogs had an AST genetic signature (24% and 28% of the total, respectively) and the average AST contents were 39% and 48%, respectively. Tests of the ability of shelter staff to classify 114 dogs >25% AST resulted in 76% correct calls; the false positive rate for 270 non-AST dogs was 1.5%. That work and previous smaller studies noted the false negative rate falls rapidly below 25-38% AST content and that the false positive rate increases with the number of breeds admixed[3,6,7]. Thus, the average such dog is approximately 50% Pit Bull and the other 50% is variable. This suggests Pit Bull-type dogs can be classified based on shelter or owner classification as we do here, but i) behavioral effects must be larger to be detected since breed effects are diluted by their breed admixture ii) it is not possible to distinguish dogs bred for fighting. Two studies from 2008 found that Pit Bull-type/mixes as defined in this study had increased rates of aggression. One included 1,448 shelter dogs classified as appearing to be, at least in part, eight breeds (Labrador, Rottweiler, Chihuahua, Husky, Chow and Beagle) or breed groups (Pit Bull and Shepherd)[8]. Standardized behavioral evaluations showed that Pit Bulls, Rottweilers, Chow Chows, Huskies, and corresponding mixes, had high risk of aggression in at least one of the nine test components (61%, 60%, 49% and 47% of dogs, respectively). The second study included 3,791 dogs with C-BARQ data and AKC-registered for 33 breeds[9]. They reported different combinations of risk of aggression traits in different breeds: Beagles and Cocker Spaniels, owner-directed; Akitas and Australian Shepherds, unfamiliar dog-directed; Chihuahuas, the previous two types and unfamiliar human-directed; Pit Bulls and Jack Russell Terriers, familiar and unfamiliar dog-directed; and Dachshund, all four types.

Dog attacks on humans have received growing public health and legal attention in the last few decades[10]. This has resulted in rules, regulations and laws that ban dogs according to their size or breed type. Despite a lack of evidence of increased human-directed aggression[9], Pit Bull-type dogs are the most likely to be banned from diverse jurisdictions in breed specific legislation. Attempts to measure the effects of such statutes showed attacks by targeted breeds were reduced, but either there was no change in overall attack rates or it was inconclusive [11-13]. The American Veterinary Society of Animal Behavior has published a position statement that claims there is insufficient evidence that breed specific legislation is effective (the legal case for this has also been made[14]) and that it has unintended negative consequences[10,15].

### Expanded results

#### Cohort

Over the last decade, analyses of canine genetics have been successful with simple traits[16,17]. However, only a small number of complex traits have been mapped[18,19]. The greatest strength of dog genetics is the large number of breeds that can be clustered into approximately ten breed groups – variation within breeds is relatively low and variation across breeds is very high. As a result, GWA within breeds or in small numbers of breeds is facilitated, but fine mapping can be difficult to impossible and carries a high risk of false positives due to population structure[20,21]. Unfortunately, linkage disequilibrium is very extensive within breeds and population structure is high (always across breeds and commonly within breeds). An additional issue is that results from single or few breeds are not representative for the full dog population.

Mixed breed dogs are a large component of the population, but are generally excluded from genetic mapping studies because they lack the power described for breeds above and have unknown population structure (that is likely complex). However, in the present study we include mixed and pure breed dogs in their existing general population proportions of roughly half and half. This is possible because our genome scans were performed in multiple cohorts with large numbers of pedigree breeds. That is, our focus was to map variants that are common across breeds. This work aims to provide further evidence for the previous interbreed findings in a cohort that represents the full community, without the need to maximize power by controlling for population structure at the level of individual breeds. Larger studies with genome wide genotyping will be necessary to further corroborate our findings while accounting for population structure.

We evaluated our cohort to estimate how representative it is of the US. All the dogs in our cohort are from US homes. 78% of the dogs are from the state of Ohio and the remainder from 26 other states. The breed makeup of pedigree dogs in our cohort is provided as Supplementary Table S1. We compared the top pedigreed breeds represented in our cohort to the most recent American Kennel Club (AKC) registration data. We used data for US breed popularity and the top five breeds in 15 cities arbitrarily selected for reporting by AKC (Suppl. Table S2). Our top 11 pedigree breeds constitute 7 ranks due to ties. Our top three breeds and ranks are identical to the top three breeds in the US and in US cities. Our top 11 breeds include US ranks 1-6, 8, 10, 13 as well as Pug (US rank 31st), Chihuahua (32nd) and Border Collie (38th). Standard Poodle was the only breed in the top 10 for the US or in the top five across US cities was absent from our cohort, but we had one Miniature Poodle and eight small Poodle mixes (presumably F1 crosses of Miniature Poodles). Given the overall breed representation, breed rankings in the US and US cities, and the fact our cohort of 397 dogs is approximately half pedigree dogs representing 77 breeds, our cohort appears to be an appropriate community sample. We suggest it can be referred to as an Ohio-biased US community sample.

We compared our cohort to others published as a point of reference. A study of inherited disorders that evaluated purebred and mixed-breed along with Pit Bull-type dogs[22] showed a higher proportion of pedigree dogs than the present study. That sample was exclusively from the University of California-Davis Veterinary Medical Teaching Hospital electronic records. Similar discrepancies are present in an epidemiological study of surgical castration[23]. This discrepancy can be attributed to the lower standard of care received by mixed breed dogs and that is associated with a lower socioeconomic status of their owners[24-26] and therefore has a lower representation in premium care facilities. This shows it is important to understand how studies differ in recruitment approach, geography and, presumably, owner socioeconomics.

**Correlations with Wilson et al. 2018 clustering of breeds by C-BARQ behavior**

Wilson et al. recently analyzed 32,005 dogs with C-BARQ data and having AKC registrations for 82 breeds (n>50 ea)[27]. They used breed means for C-BARQ behavioral subscales to calculate an Euclidian distance matrix. Agglomerative hierarchical clustering performed on that matrix resulted in all breeds being assigned to six clusters (no significance or confidence measures were given). One cluster included the American Staffordshire Terrier (AST) and all of the breeds known to be most closely related to it: Boxer, Bulldog, French Bulldog, Bull Terrier and Boston Terrier, and two unexpected, Dalmatian and Italian Greyhound. Members of several other breed groups also tended to be enriched in one or two clusters. However, the contribution of breed group to the clustering was secondary to the mean height of breeds. For instance, there is a cluster of mostly ancient/Asian breeds, but other members of that group are in a large group of the shortest breeds. The shepherd group breeds largely lie in a cluster of mostly shepherds and retrievers, but the Pembroke Welsh Corgi is in the cluster enriched for terriers and the Collie is clustered with all of the tallest breeds. The top level finding of the study was the correlation of body size and the C-BARQ behavior matrix. Across most behaviors, the increased risk of problem behaviors (and non-trainability) tends to be correlated with decreased height. The AST cluster means are not the most extreme positively or negatively for any C-BARQ behavioral subscale. In contrast to the clusters with smallest and largest average body size, the AST cluster is within one standard deviation of the mean of all clusters for all C-BARQ behavioral subscales as well as miscellaneous traits. The two most notable subscale effects are slightly reduced risk of aggression directed at unfamiliar humans and slightly increased energy. The AST cluster had the most extreme mean scores for chewing of inappropriate objects, hyperactivity, “chasing shadows” and, marginally, staring.

Given that the Wilson et al. study[27] was able to cluster according to C-BARQ behavioral data, we compared that clustering to our present findings. We classified all of our pedigree and Pit Bull-type dogs according to Wilson et al.’s clusters (Figs. S13, S14). The PCA of genetic markers in our cohort shows Wilson et al.’s clusters partially segregate in our data (Fig. S13). The clearest cluster is AST, which is the only one almost entirely in the left bottom quadrant (i.e., negative in PC1 and 2). All members of that Wilson et al. cluster in our cohort are breeds related to Pit Bull-type dogs, except for three Italian Greyhounds (i.e., one of the genetically-unexpected members of the group because they are not closely related to Pit Bull-type dogs). The latter are the only three members of this cluster >1 in PC1, and all are approximately 2-3 in PC1. Only one of Wilson et al.’s breed groups is in that range or higher in PC1: the “Maltese” cluster which is most distinct for breeds of small body size. This is consistent with Italian Greyhounds, which are a miniature breed, grouping by genetic markers with small dogs. Other Wilson et al. clusters are evident in our data, but less distinct (e.g., shepherd, ancient/Asian, and smallest and largest breed clusters). Overall, the observed patterns for the genetic marker PCA are consistent with

Wilson et al.'s cluster associations with height and breed relatedness. The Pit Bull-type dog findings support their classification in this study. The distribution of the Wilson et al. AST cluster dogs in Figure S9 and the tighter grouping of a subset of Pit Bull-type dogs in the far left and bottom in Figure 2C suggest the latter are the most homogenous and, therefore, purest Pit Bull-type dogs (i.e., have the highest AST content[4]).

Thus the Wilson et al.[27] behaviorally-defined clusters of breeds grossly segregate in the PCA of 20 genetic markers at 13 loci in our weakly powered study (n=400 vs. Wilson et al.'s n=32,000). We presume this is primarily due to genetic variation associated with body size and breed-relatedness. In contrast, PCA of our behavioral data (Fig. S14) did not yield any clear grouping of dogs according to the Wilson et al. clusters. This shows that the associations between our genetic markers and behaviors cannot be explained by correlations of breeds and behaviors according to genetic relatedness (i.e., due to both breed phylogenetics and body size admixture). That is consistent with Wilson et al.'s main findings: breeds clustered by behavior have some effect of breed relatedness, but breed height is a better predictor and some breed groups have members in many behavioral clusters.

### Expanded discussion

**Summary of the present results in context of interbreed behavioral genome scans**

Our original GWAS's of fear and aggression traits yielded 16 significant loci, four of which were associated with the same 10 traits in two cohorts[28]. Here we provide further evidence for all four of those loci and most of the traits, either directly or as related traits. Chr10 was further supported in this study for separation anxiety (in the FMM), but not for touch sensitivity. We previously reported that chr15 was associated with owner-directed aggression in two cohorts as well as familiar dog aggression, unfamiliar dog-directed fear, separation anxiety and touch sensitivity in a single cohort. Here we found chr15 is associated with dog-directed aggression, touch sensitivity, non-social fear, separation urination and barking in the FMM, and some of those plus many other, mostly anxiety or separation-related, traits in the IMM. However, neither model was significant for owner-directed aggression. Assuming that the original finding was true, an interesting possibility is that genetic modifier variations that increase the risk of this phenotype are under negative selection or diluted by mixed breeding. Because the chr15 allele is fixed in many breeds, it could only be under negative selection in the other breeds or in mix-breed dogs (chr15/*IGF1* and size are discussed below). Another possibility is that participating owners in the present study are somehow different in comparison to prior studies. For example, we could have fewer dogs with owner-directed aggression here due to increased relinquishment/euthanasia in the shelters or reduced owner participation. Chr18 was originally mapped in two cohorts for three fear or aggression traits directed at unfamiliar dogs and humans (and in one cohort for the fourth of those), but not aggression directed at familiar humans or dogs; and touch sensitivity and non-social fear. Of those, we provide further evidence for aggression directed at unfamiliar humans and dogs in both models, as well as fear of unfamiliar dogs in the IMM. Both models also showed chr18 is associated with increased separation urination and barking, as well as decreased leash pulling. We originally reported chrX was associated with dog-directed fear and separation anxiety in two cohorts, as well as other social and non-social fear and anxiety traits, but not aggression directed at familiar humans or dogs. Here we provide further evidence that unfamiliar dog-directed fear and found other chrX associations with submissive/emotional urination, separation defecation and rolling in odors (feces and other highly odorous substances).

Other marker-trait associations that support the genome scans include the following: chr1B and dog-directed fear (IMM; each study also had a different separation-related association); chr10A (a very distant locus from chr10B-E) and unfamiliar dog-directed aggression (both models); chr20 and chasing (both models); and chr34 with aggression and fear. More specifically, chr34 was mapped for unfamiliar dog-directed fear and aggression, excitability and escaping, but here was associated with unfamiliar and familiar human-directed fear, non-social fear, touch sensitivity, submissive emotional urination, separation defecation, fear of stairs and “other behaviors”. Chr32 was mapped for familiar dog-directed aggression and excitability and here is associated with unfamiliar human-directed fear and aggression, unfamiliar dog-directed aggression, attention seeking, separation anxiety and touch sensitivity. Chr32 is negatively associated with fear of stairs, but is not directly associated with body size (a trait known to be negatively correlated with fear of stairs[29]). Chr32 is the only marker associated with fear of stairs in the FMM, positively or negatively. In contrast, only one marker known to be associated with size is also associated with fear of stairs, chr15, but that is only in the IMM. [Note on chr32 and body size: Although the chromosome 32 marker is correlated with known size markers on 10, 15 and X, the relationship between size and the chr32-association with fear of stairs correlates with increased size for chr10 and chrX, and decreased size for chr15. Comparison of the two markers for FMM shows 6 trait associations for chr15 (small size allele) and two for chr32, and no overlap between them. For IMM, there are 12 trait associations with chr15 and 7 with chr32, and two overlap – one in the same direction and one in the opposite direction (the latter is fear of stairs).] Chr1A was mapped for excitability but here was associated with non-trainability, which overlaps with excitability in the PCA (Fig. 2B). Chr13 had different, but similar or related, social fear and aggression associations in the mapping discovery and present studies. Chr24A and B are close to each other, but have both shared and non-shared association types in the genome scans: they shared anxiety or separation associations, but only 24B was associated with chasing and energy. However, here we found 24A is associated with hyperactivity and 24B with rolling in odors. This suggests that, if there are two chr24 variants, they are frequently carried together.

**On the possibility of pleiotropy**

There is growing evidence of extensive pleiotropy in human complex genetics[30,31]. Watanabe et al. performed joint analyses of reasonably powered (N > 50,000) GWAS’s to date, which represent 558 unique traits across 24 trait domains. Based on that, they generated an overview of pleiotropy in complex traits. They found gene-level pleiotropy for 81% of genes associated with any trait, 67% of which were multidomain. Variant or SNP-level pleiotropy was observed for 60% of trait associations, 32% of which were multidomain. Their supplementary information included the gene pleiotropy data. Of the top candidate gene at each of the 13 loci in our present study, 12 were present (Suppl. Table S15). 10 of those 12 showed multidomain pleiotropy; of the other two, one was psychiatric and one immunological. Of the 12, seven included at least one of the domains most relevant to behavior: Activities, Psychiatric or Social Interactions. Of the latter seven, six were multidomain and five included one of the two domains most relevant to body size and morphology: Metabolism and Skeletal. This shows that, at the gene level alone, plausibility for our candidates to have morphological and behavioral pleiotropy is strong.

**Canid biology and human interactions**

Barking is one example of the intersection of dog biology and human environmental effects on their behavior. A prominent difference between wolves and dogs is barking behavior. Whereas wolves bark to warn and protest in social contexts, domesticated dogs have an earlier onset, lower threshold and broader set of contexts and purposes[32]. Barking can be a highly desirable dog trait for humans, such as for warning of danger or uncommon circumstances and intimidation of adversaries and predators. It seems likely that barking was selected in early domestication, but it cannot be ruled out that selection for an associated trait is responsible. Today, dog barking is one of the most common human complaints in the behavioral clinic and at large. However, it is not difficult to imagine there could be major life course behavior differences in genetically identical dogs that are rewarded vs. punished for barking. Here we showed positive association of barking with the presence of other animals in the home, behavioral diagnosis and behavioral medication, but negative association with non-behavioral medical conditions. The barking-correlated loci were chr15 (both models), chr18 and chr20. Future studies can isolate each of those loci and measure association with novelty and fear stimuli, penetrance, sexual sterilization, environmental effects (incl. human behavior) and therapeutic response.

A more mysterious dog trait, coprophagia, has a prevalence of 16% for common occurrence and 24% for any, but no clear purpose[33]. Wolf mothers are known to eat their pups' stools, which is assumed to represent cleaning the den. Some have proposed domesticated dogs may do the same for similar purposes[33]. Others have suggested it is part of a normal canid scavenging behaviors, consistent with consumption of stools from diverse species (incl. horse, cattle, deer, rabbit, cat and human) in addition to a reduced rate for canine and their own[34]. Among the purposes of coprophagia allegedly ruled out have been nutritional deficiency, gastrointestinal disease and housetraining difficulties (suggestive of deficiency in feces aversion). In order of increasing p-value, the traits previously known to be associated with coprophagia are greedy eating, eating dirt, multiple-dogs in household, and breed group[33]. Chasing away from stools, punishing with electronic and sound-emitting collars, anti-coprophagia-purposed food additives/tablets and pepper-lacing of stools had 0-2% efficacy in preventing coprophagia; reward of the successful command of "leave it alone" was 4%. Here we found that coprophagia is associated with non-behavioral medical conditions, the presence of other animals in the home, behavioral diagnosis and the loci on chr10 and 20. Curiously, both of the latter associations are in the opposite direction of the majority of other associations with the same loci – most of which are related to aggression and anxiety. A rapidly growing area of interest is the modulation of behaviors such as aggression, anxiety and depression by the gut microbiome and its influence on neuroendocrine and immune systems[35,36]. Future studies would be necessary to consider if coprophagia could have probiotic or therapeutic effects on inflammation or behavior, including anxiolytic or anti-aggressive effects. This seems consistent with the observation that canine coprophagia involves fresh feces (up to 2 days old) and 85% of the source is non-self[33]. Some of the coprophagia associations in this study also support this possibility, mainly those of behavioral and non-behavioral medical diagnoses (Figs. 4 and 5).

Other associations are additional animals in the home, being a working dog, being a competition dog, and the chr10E and chr20 markers. Being a Pit Bull-type dog and chr1A are protective (Figs. 4 and 5;

Suppl. Fig. S9). The traits most strongly correlated with coprophagia were rolling in odors, chewing inappropriate objects, and grooming others. Several other traits are also correlated though less strongly (Suppl. Fig. S2). Importantly, none of the fear or aggression traits or non-social fear were associated with fecal-ingestion. These observations indicate coprophagia is negatively associated with behavioral and non-behavioral wellbeing, but is not related to fear.

In this cohort the presence of children in the home was associated with a decreased incidence of behavioral diagnoses. Child contributions to this could be complex, but the effect may be related to parental diligence to protect children by removing dogs with problem behaviors. A study of behaviorally diagnosed dogs showed that the proportion of anxiety/separation vs. aggression diagnoses is highest for single owner homes, then couples and, lastly those with children (75, 65 and 54 percent anxiety, respectively[37]). Children in the home could have an overall effect on mental state that reduces some aspects of anxiety but possibly increases aggression (presumably due to fear). Here only two traits were positively associated with children in the home in both modeling modes: snapping at flies and shadow chasing. Other associations with children were increased separation urination (FMM), chewing on inappropriate objects and stealing food (IMM), and decreased "other behaviors" (FMM) and unfamiliar human-directed fear (IMM). It will be important to determine if stealing food could somehow be related to food guarding behaviors. Whereas the most common circumstance for bites to unfamiliar children is territorial guarding, the most common for familiar children is food guarding[38].

The strongest associations with the presence of children in the home in our cohort were with snapping at flies and chasing shadows. Since stereotypic or compulsive behaviors like these are commonly indicative of stress, this is consistent with canine stress being elevated in households with children. However, those two traits are not the most likely to result in clinical assessment. Although one model each showed association of children and increased separation urination and inappropriate chewing, there was also reduced risk of unfamiliar human-directed fear and "other behaviors". In the IMM, which allows the most direct isolation of individual effects, the four associations that increase risk (inappropriate chewing, stealing food, snapping at flies and chasing shadows) were unique to children in the home out of the 17 questionnaire predictor variables. This suggests the observed effects are not due to correlations with other predictor variables (i.e., and their association with trait variables). Stealing food seems likely to be, at least in part, a learned behavior. In homes where children drop or throw food around there is great temptation for dogs to "steal" food, and similar effects seem possible for inappropriate chewing. PC2 placed inappropriate chewing and unfamiliar human-directed fear at the opposite extremes (Fig. 2B). This is interesting because children in the home increased risk for the first of those traits but is protective against the second. Stealing food is the second most extreme at the chewing of inappropriate objects end, and chasing shadows and snapping at flies are also in the top third of the behaviors with increased risk associated with the presence of children in the home. Separation urination appeared slightly below snapping at flies. This is consistent with children having a particular impact on a biological phenomenon related to PC2. The large effect of having children in the home on traits associated with our present risk loci suggest this may be an ideal area in which to identify gene-environment interactions.

**Pit Bull-type dog behavior**

Approximately 15% of our cohort is Pit Bull-type according to owner reporting. The next highest frequency was 6% for all retrievers. Pit Bull-type in our study has some phenotypic uniformity, but not as strong as a breed (Fig. 2C and Suppl. Fig. S5). This is expected because it is presumably enriched for one breed type, but frequently mixed with diverse other breeds (here 51/59 were reported to be mixed breed). Isolation of Wilson et al. behavioral clusters in the PCA of our genetic markers shows strong grouping of related breeds (see last section of Suppl. Text, Expanded results; Suppl. Fig. S13). Behaviorally, Pit Bull-type dogs are relatively heterogeneous in comparison with the rest of our cohort. As a group, they were not associated with behavioral diagnosis (Fig. 1). Our PCA of genetic markers shows a subset of Pit Bull-type dogs cluster together (Fig. 2C), but PCA of behavioral traits lacks such an effect (Fig. 2D). That means that the variants captured by the 20 markers of this study do not segregate Pit Bull-type dogs into behavioral subgroups.

Only one problem trait is associated with Pit Bull-type in the generalized linear models, pulling on the leash (FMM and IMM; Figs. 4, 5). Threshold logistic regression with stepwise selection modeling revealed Pit Bull-type dogs have reduced owner directed aggression at the 75$^{th}$ quantile threshold and increased unfamiliar dog-directed fear at the 95$^{th}$ (Suppl. Figs. S10 and S12). Consistent with the first of those, a large study of aggression using C-BARQ data for dogs from 32 AKC breeds and Pit Bull-type dogs (as defined here) showed the latter had significantly reduced levels of owner-directed aggression[9]. They reported both breed means and the percent of breed members with the maximum score for any of four aggression types. Those data indicated that a subset of at least 11.5% of Pit Bull-type dogs have extreme aggression (significantly higher than the breed mean) directed at familiar dogs. This suggests the possibility that the 95$^{th}$ quantile Pit Bull-type enrichment we observe for fear of unfamiliar dogs could be related to that. This does not seem surprising to us with respect to the correlation of fear and aggression. However, whereas there is a great distinction between aggression directed at familiar vs. unfamiliar humans, the issue of familiarity is less clear for dog-directed aggression. While the previous study analyzed aggression but not fear traits, we analyzed both. If we assume Pit Bull-type dog behavior and C-BARQ reporting were similar in the two studies, our findings would suggest owners detect fear and aggression differently. Specifically, aggression could be perceived more sensitively compared to fear (either in absolute terms or due to scaling of severity). The present study is underpowered to establish whether the extreme fear and extreme aggression Pit Bull-type dogs are the same, but we predict this will be proven to be the case.

We did not find evidence that Pit Bull-type dogs are exceptionally aggressive in our cohort. One important consideration is that the Pit Bull-type dogs in our cohort may not be representative of the full population. For instance, the most successful fighting dogs seem less likely to be relinquished to shelters and would be under-represented in our cohort. We also cannot rule out the possibility that Pit Bull-type dogs may have unique variations that are associated with aggression, but are rare in most breeds. Such variation would not detectable in our cross-breed mapping studies and would not be included in the markers used in this study. Top markers shared by Pit Bull and non-Pit Bull-type dogs that should be prioritized for further behavioral and physiological investigation include chr10C, associated with aggression directed at unfamiliar humans, and chr13 and chrXB, which are associated with dog-directed fear. Based on our prior study, chr18[(28)] should be evaluated as well for its association with dog-directed fear. We predict that co-inheritance of those variants is likely to result in increased risk and severity of fear/aggression phenotypes regardless of breed status[28].

It is important to further define the genetics of Pit Bull-type dog behavior, including fear and aggression. Such studies could include more specific definitions of the leash pulling trait we found (incl. by association with other behaviors). An interesting possibility is that the behaviors selected in the ancestral fighting line were not simply related to increased overall aggression. Diane Jessup, a US Pit Bull aficionado with long term success in Schutzhund and other types of competitions, contended that the fighting bloodline (aka, game or game-bred dogs) has desirable non-aggression traits such as work drive and trainability[5]. In contrast the Staffordshire Bull Terrier, the extant parental-breed in the UK, has allegedly been selected for docility since dog fighting was made illegal in the UK in 1835[5]. It would not be surprising if the ancestral fighting line was selected for traits such as physical strength and stamina, boldness, fearlessness, work drive and human-controlled behavior including but not limited to fighting. Even if that was once the norm, it seems likely that such fighting-purposed dogs were mixed with dogs exhibiting increased fear-related aggression. Only at the 95$^{th}$ percentile of severity in threshold modeling, was unfamiliar dog-directed fear associated with Pit Bull-type dogs. It would be interesting to evaluate whether that fear variation, which is rare but had a large effect, is part of the early Pit Bull genome or was introduced more recently. Regardless of the selection history, this suggests it may be possible to identify and remove such variation from the Pit Bull. At the same time, it will be important to determine if any of those variations are associated with other beneficial traits. Aside from public health and statutory issues, these studies could provide new insights on the biology of aggression.

### Table S1. Dog breed frequencies included in the study

| Breed | BreedID | Frequency | Percent | Cumulative Percent |
|---|---|---|---|---|
| American Bulldog | ABulldog | 1 | 0.25 | 0.25 |
| American Eskimo Dog | AEskimoD | 1 | 0.25 | 0.5 |
| Akita | Akita | 2 | 0.5 | 1.01 |
| Australian Cattle Dog | AusCattl | 3 | 0.76 | 1.76 |
| Australian Shepherd | AusShep | 4 | 1.01 | 2.77 |
| Australian Shepherd Toy | AusShepT | 1 | 0.25 | 3.02 |
| Basset Hound | BassetHo | 2 | 0.5 | 3.53 |
| Beagle | Beagle | 6 | 1.51 | 5.04 |
| Belgian Groenendael | BelgGroe | 1 | 0.25 | 5.29 |
| Belgian Malinois | BelgMali | 2 | 0.5 | 5.79 |
| Bernese Mountain Dog | BernesMD | 1 | 0.25 | 6.05 |
| Bishon Frise | BichonFr | 3 | 0.76 | 6.8 |
| Black Mouth Cur | BlackMoC | 1 | 0.25 | 7.05 |
| Border Collie | BordColl | 8 | 2.02 | 9.07 |
| Boston Terrier | BostonTe | 3 | 0.76 | 9.82 |
| Boxer | Boxer | 7 | 1.76 | 11.59 |
| Bulldog (unpecified) | Bulldog | 2 | 0.5 | 12.09 |
| Curly Coat Retriever | CCoaRet | 1 | 0.25 | 12.34 |
| Cavalier King Charles Spaniel | CKCharle | 1 | 0.25 | 12.59 |
| Cane Corso | CaneCors | 1 | 0.25 | 12.85 |
| Carolina Dog | Carolina | 1 | 0.25 | 13.1 |
| Chihuahua | Chihuahu | 5 | 1.26 | 14.36 |
| Chow Chow | ChowChow | 1 | 0.25 | 14.61 |
| Cockapoo | Cockapoo | 3 | 0.76 | 15.37 |
| Collie | Collie | 2 | 0.5 | 15.87 |
| Dachshund | Dachshun | 5 | 1.26 | 17.13 |
| Doberman Pinscher | Doberman | 2 | 0.5 | 17.63 |
| English Bulldog | EBulldog | 4 | 1.01 | 18.64 |
| English Shepherd | EnglishS | 1 | 0.25 | 18.89 |
| Fox Terrier | FoxTerri | 4 | 1.01 | 19.9 |
| French Bulldog | FrenchBu | 1 | 0.25 | 20.15 |
| Great Pyrenees | GPyrenee | 3 | 0.76 | 20.91 |
| German Shepherd | GerShep | 11 | 2.77 | 23.68 |
| German Wirehaired Pointer | GerWireP | 1 | 0.25 | 23.93 |
| Golden Retriever | GoldRet | 10 | 2.52 | 26.45 |
| Great Dane | GreatDan | 7 | 1.76 | 28.21 |
| Greyhound | Greyhoun | 2 | 0.5 | 28.72 |
| Havanese | Havanese | 3 | 0.76 | 29.47 |
| Italian Greyhound | ItalianG | 3 | 0.76 | 30.23 |
| Jack Russel Terrier | JackRuss | 1 | 0.25 | 30.48 |
| Kangal Dog | KangalDo | 1 | 0.25 | 30.73 |
| Labrador Retriever | LabRet | 13 | 3.27 | 34.01 |
| Large Munsterlander | Lmunster | 1 | 0.25 | 34.26 |
| English Mastiff | Mastiff | 1 | 0.25 | 34.51 |
| Miniature Pinscher | MinPinsc | 1 | 0.25 | 34.76 |
| Mixed Breed | Mixed | 188 | 47.36 | 82.12 |
| Olde English Bulldog | OEBulldo | 1 | 0.25 | 82.37 |
| Old English Shepherd | OEngShee | 1 | 0.25 | 82.62 |
| Papillon | Papillon | 1 | 0.25 | 82.87 |
| Parson Russel Terrier | ParsonRu | 1 | 0.25 | 83.12 |
| Pitbull (thus reported; not incl. mix) | Pitbull | 18 | 4.53 | 87.66 |

| | | | | |
|---|---|---|---|---|
| Pointer | Pointer | 2 | 0.5 | 88.16 |
| Pomeranian | Pomerani | 2 | 0.5 | 88.66 |
| French Poodle Miniature | PoodleMi | 1 | 0.25 | 88.92 |
| French Poodle Toy | PoodleTo | 1 | 0.25 | 89.17 |
| Portuguese Water Dog | PortugWD | 2 | 0.5 | 89.67 |
| Pug | Pug | 5 | 1.26 | 90.93 |
| Rat Terrier | RatTerri | 1 | 0.25 | 91.18 |
| Redbone Coonhound | RedCoonh | 1 | 0.25 | 91.44 |
| Rhodesian Ridgeback | Rhodesia | 1 | 0.25 | 91.69 |
| Rottweiler | Rottweil | 8 | 2.02 | 93.7 |
| Soft Coal Wheaten Terrier | SCWheate | 1 | 0.25 | 93.95 |
| Saint Bernard | SaintBer | 1 | 0.25 | 94.21 |
| Schnauzer Medium | SchnauzM | 1 | 0.25 | 94.46 |
| Schnauzer Small | SchnauzS | 1 | 0.25 | 94.71 |
| Scottish Deerhound | ScottDee | 1 | 0.25 | 94.96 |
| Shiba Inu | ShibaInu | 2 | 0.5 | 95.47 |
| Shih Tzu | ShihTzu | 1 | 0.25 | 95.72 |
| Shih-poo | Shihpoo | 3 | 0.76 | 96.47 |
| Siberian Husky | SibHusky | 2 | 0.5 | 96.98 |
| Springer Spaniel | Springer | 1 | 0.25 | 97.23 |
| Staffordshire Bull Terrier | Stafford | 3 | 0.76 | 97.98 |
| Yorkshire Terrier Teacup | TeacYork | 1 | 0.25 | 98.24 |
| Treeing Walker Coonhound | TrWalkCo | 1 | 0.25 | 98.49 |
| Whippet | Whippet | 2 | 0.5 | 98.99 |
| Wirehaired Pointing Griffon | WiPoinGr | 1 | 0.25 | 99.24 |
| Yorkipoo | Yorkipoo | 1 | 0.25 | 99.5 |
| Yorkshire Terrier | Yorkshir | 2 | 0.5 | 100 |
| TOTAL | | 397 | | |
| 77 distinct Purebreeds + Mixed Breed | | | | |

### Table S2. Comparison of cohort breeds to popularity in US and US cities

| | Top 11 Breeds in Present Cohort Rank (# dogs) | US Popularity Rank[1] | 15 Cities Popularity Rank (# cities in top 5)[1] |
|---|---|---|---|
| Labrador Retriever | 1 (13) | 1 | 1 (15) |
| German Shepherd Dog | 2 (11) | 2 | 2 (13) |
| Golden Retriever | 3 (10) | 3 | 1 (15) |
| Rottweiler | 4 (8) | 8 | 6 (1) |
| Border Collie | 4 (8) | 38 | (none) |
| Boxer | 5 (7) | 10 | (none) |
| Great Dane | 5 (7) | 14 | (none) |
| Bulldog | 6 (6) | 5 | 3 (10) |
| Beagle | 6 (6) | 6 | 6 (1) |
| Chihuahua | 7 (5) | 32 | (none) |
| Dachshund | 7 (5) | 13 | (none) |
| Pug | 7 (5) | 31 | (none) |
| French bulldog | (1) | 4 | 2 (13) |
| Yorkshire Terrier | (2) | 9 | 4 (3) |
| [2]*Standard Poodle* | *(0)* | *7* | *5 (2)* |
| [3]*Pembroke Welsh Corgi* | *(0)* | *15* | *5 (2)* |

[1]Source: American Kennel Club (2017 US data; 2018 15 arbitrary cities data: Atlanta, Miami, Chicago, Dallas, New York, Houston, Los Angeles, Boston, Denver, Philadelphia, Raleigh, North Carolina, San Francisco, Seattle, Washington, D.C., West Palm Beach, Florida)
[2]Standard Poodle pedigree breed is absent, but we have 1 Miniature Poodle and 8 small Poodle mix (presumably, F1 crosses of Miniature Poodles)
[3]Pembroke Welsh corgi popularity suggests regional effect (only ranked in San Francisco and Seattle)

**Table S3. Markers list with genome scan traits**

| Present nomenclature | Marker ID | canfam3 assembly | Mapped behavioral traits | Nearest genes[3] (distance, kb) | Gene (brief function[4]) | Zapata et al. 2016 GWA locus | No. cohorts with locus association[5] |
|---|---|---|---|---|---|---|---|
| chr1A | BICF2P1070568 | chr1:42,035,784 | Excitability | ***ESR1*** | Estrogen Receptor 1 (ligand-activated transcription factor) | n | 1 |
| chr1B | BICF2P573696 | chr1:93,250,465 | SeparationAnxiety, DogFear, Chasing | *RCL1* (46.1) | RNA Terminal Phosphate Cyclase Like 1 (has roles in ribosome biogenesis and rRNA processing) | n | 2 |
| chr5 | BICF2P1259166 | chr5:35635596 | Escaping, Chasing | ***SHISA6*** | Shisa Family Member 6 (regulates AMPA-type glutamate receptor (AMPAR) immobilization at postsynaptic density keeping the channels in an activated state in the presence of glutamate and preventing synaptic depression) | n | 1 |
| chr10A | BICF2S2363180 | chr10:2,431,382 | TouchSensitivity, Excitability, DogAggression, TouchSensitivity, NonsocialFear, Attachment, SeparationUrination, StrangerAggression | *LRIG3* (290.0) | Leucine Rich Repeats And Immunoglobulin Like Domains 3 (known to have roles in morphogenesis) | y | 2 |

| | | | | | | | |
|---|---|---|---|---|---|---|---|
| chr10B | BICF2P283272 | chr10:7,996,770 | [1]Excitability, Attachment, OwnerAggression, TouchSensitivity, SeparationUrination, SeparationAnxiety, Barking | ***MSRB3*** | Methionine Sulfoxide Reductase B3 (catalyzes the reduction of methionine sulfoxide to methionine) | y | 3 |
| chr10C | BICF2S23012887 | chr10:8,059,173 | [1]Excitability, SeparationAnxiety, Escaping | *MSRB3* (18.0) | " | y | 3 |
| chr10D | BICF2P709352 | chr10:8,397,696 | [1]RivalryAggression | ***HMGA2*** | High Mobility Group AT-Hook 2 (non-histone chromosomal protein with functions of architectural factor and essential component of the enhancesome) | y | 3 |
| chr10E | BICF2P776298 | chr10:8,454,499 | [1]Attachment, Excitability, DogAggression, Excitability, Attachment, SeparationUrination, Barking | ***HMGA2*** | " | y | 3 |
| chr13 | BICF2P1389702 | chr13:8,391,652 | Excitability, SeparationAnxiety, Attachment, DogAggression, NonsocialFear | *ANGPT1* (79.5) | Angiopoietin 1 (secreted glycoprotein with roles in vascular development and angiogenesis) | n | 2 |
| chr15A | BICF2P885652 | chr15:41,232,547 | SeparationUrination, OwnerAggression, TouchSensitivity, OwnerAggression | ***IGF1*** | Insulin Like Growth Factor 1 | y | 3 |
| chr15B | BICF2P117496 | chr15:41,250,986 | TouchSensitivity, RivalryAggression, SeparationUrination | ***IGF1*** | " | y | 3 |

| | | | | | | | |
|---|---|---|---|---|---|---|---|
| chr18 | BICF2S2323033 | chr18:20,310,833 | StrangerFear, DogFear, NonsocialFear, StrangerAggression, TouchSensitivity, OwnerAggression, Barking, Trainability, SeparationAnxiety, Attachment, DogAggression, RivalryAggression, Chasing, SeparationUrination, Excitability | ***CD36*** | CD36 (multifunctional glycoprotein that acts as receptor for a broad range of protein and lipid ligands) | y | 3 |
| chr20 | BICF2G630233714 | chr20:21,848,178 | Escaping, Chasing, OwnerAggression, SeparationUrination, SeparationAnxiety | ***MITF*** | Melanocyte Inducing Transcription Factor (transcription factor that regulates melanocyte development and is responsible for pigment cell-specific transcription of the melanogenesis enzyme genes) | n | 3 |
| chr24A | TIGRP2P314845 | chr24:23,196,435 | NonsocialFear, TouchSensitivity, RivalryAggression, SeparationAnxiety, OwnerAggression | ***RALY*** | Marker may be in same haplotype as chr24B at least in some breeds: RALY Heterogeneous Nuclear Ribonucleoprotein (aka, HnRNP-Associated With Lethal Yellow; roles in pre-mRNA splicing, embryonic development | n | 2 |
| chr24B | BICF2P131040 | chr24:23,398,090 | SeparationUrination, Energy, Chasing | *ASIP* (4.0) | Agouti Signaling Protein (paracrine signaling molecule involved in melanogenesis and pigmentation color) | n | 2 |

| | | | | | | | |
|---|---|---|---|---|---|---|---|
| chr32 | BICF2G630600200 | chr32:5,421,210 | Excitability, FamDogAggr | *PRKG2* (55.8); *RASGEF1B* (123.5) | Protein Kinase CGMP-Dependent 2 (serine/threonine protein kinase with roles in intestinal secretion and bone growth); RasGEF Domain Family Member 1B (Guanine nucleotide exchange factor) | y | 1 |
| chr34 | BICF2P183789 | chr34:18,559,537 | Excitability, Escaping, DogAggression, DogFear | *IGF2BP2* (37.4) | Insulin Like Growth Factor 2 MRNA Binding Protein 2 (binds insulin-like growth factor 2 mRNA and regulates its translation) | y | 1 |
| chrXA | BICF2P614144 | chrX:101,646,292 | [1]Chasing | *ENSCAFG00000018773* (10.9); *ENOX2* (96.1); [2]<u>*IGSF1*</u> | ENSCAFG00000018773 (no annotation)[2]; Ecto-NOX Disulfide-Thiol Exchanger 2 (cell surface NADH oxidase with two enzymatic activities: catalysis of hydroquinone or NADH oxidation, and protein disulfide interchange) | y | 2 |
| chrXB | BICF2S23759452 | chrX:102,553,876 | [1]DogFear, Chasing | [2]<u>*IGSF1*</u> (180.0) | Immunoglobulin Superfamily Member 1 (participates in the regulation of interactions between cells) | y | 2 |
| chrXC | BICF2P350270 | chrX:103,129,523 | [1]Barking, DogFear, SeparationAnxiety | *RAP2C* (5.1); [2]<u>*IGSF1*</u> | RAP2C (Ras-related protein subfamily of the Ras GTPase superfamily that regulate cell proliferation, differentiation and apoptosis)[2] | y | 2 |

[1]For these multiple markers at one locus, only refers to traits associated with that marker rather than all markers in the locus
[2]IGSF1 implicated to be the causative gene variant for size and fear at this locus with extensive linkage disequilibrium (PMID: 28257443; Zapata et al. 2016)
[3]SNP markers within genes in bold
[4]Source: GeneCards
[5]Loci mapped in one cohort were selected for the biochemistry or biological relevance of candidate genes

### Table S4. Descriptive statistics for continuous variables

| Variable | Definition | N | Mean | Std Dev |
|---|---|---|---|---|
| Weight | Weight in lbs | 394 | 22.954 | 14.214 |
| Age when acquired (AcquirAGE) | Age in weeks when acquired | 390 | 59.908 | 95.338 |
| Age when evaluated (AgeAtEvaluation) | Age in weeks when evaluated | 397 | 277.700 | 192.619 |
| Trainability (Train) | Willingness to attend to the owner, obey simple commands, fetch objects, respond positively to correction, and ignore distracting stimuli | 393 | 2.518 | 0.674 |
| Stranger-directed aggression (StrDirAgg) | Threatening or aggressive responses to strangers approaching or invading the dog's or owner's personal space, territory, or home range | 387 | 0.989 | 0.946 |
| Owner-directed aggression (OwnDirAgg) | Threatening or aggressive responses to the owner or other members of the household when challenged, manhandled, stared at, stepped over, or when approached while in possession of food or objects | 397 | 0.234 | 0.437 |
| Dog-directed aggression (DogDirAgg) | Threatening or aggressive responses when approached directly by unfamiliar dogs | 367 | 1.458 | 1.219 |
| Familiar dog aggression (FamDogAgg) | Aggressive or threatening responses to other familiar dogs in the household | 329 | 0.762 | 1.007 |
| Dog-directed fear (DogDirFear | Fearful or wary responses when approached directly by unfamiliar dogs | 371 | 1.118 | 1.122 |
| Stranger-directed fear (StrDirFear) | Fearful or wary responses when approached directly by strange or unfamiliar people | 391 | 0.971 | 1.187 |
| Nonsocial fear (NonSocFear) | Fearful or wary responses to sudden or loud noises, traffic, and unfamiliar objects and situations | 390 | 1.164 | 0.953 |
| Touch Sensitivity (TouchSen) | Fearful or wary responses to potentially painful or uncomfortable procedures, including bathing, grooming, nail-clipping, and veterinary examinations | 390 | 1.057 | 0.959 |
| Separation-related problems (SepRelProb) | Vocalizing and/or destructive behavior when separated from the owner, including autonomic signs of anxiety—restlessness, loss of appetite, trembling, and excessive salivation | 394 | 0.795 | 0.825 |
| Excitability (Excite) | Reaction to potentially exciting or arousing events, such as going for walks or car trips, doorbells, arrival of visitors, or the owner arriving home; difficulty settling down after such events | 394 | 2.450 | 0.834 |
| Attachment/Attention-seeking (AtcAtnSeek) | Maintains close proximity to the owner or other members of the household, solicits affection or attention, becomes agitated when the owner gives attention to third parties. | 396 | 2.246 | 0.852 |

| | | | | |
|---|---|---|---|---|
| Chasing | Pursues cats, birds, and/or other small animals, given the opportunity | 390 | 2.412 | 1.074 |
| Energy level (Energy) | Level of energetic, boisterous, and/or playful behavior | 397 | 1.987 | 1.158 |
| Escaping and roaming (EscapeRoam) | Escapes and roams from home or yard, given the opportunity. | 378 | 1.519 | 1.380 |
| Rolling in odors (Rolling) | Rolls in animal droppings or other strong odors | 391 | 1.307 | 1.356 |
| Coprophagia | Eats own or other animal's droppings or feces | 392 | 0.954 | 1.289 |
| Chewing | Chews inappropriate objects | 396 | 1.010 | 1.189 |
| Mounting | Mounts or 'humps' objects, furniture or people | 394 | 0.416 | 0.856 |
| Begging | Begs persistently for food when people are eating | 396 | 1.768 | 1.349 |
| Food Stealing (FoodSteal) | Steals food | 393 | 1.097 | 1.250 |
| Fear or stairs (FearStairs) | Nervous or frightened on stairs | 389 | 0.416 | 0.906 |
| Leash pulling (PullLeash) | Pulls excessively hard when on leash | 395 | 1.554 | 1.276 |
| Urine marking in home (MarkUrine) | Urinates against objects/furnishings in the home | 396 | 0.227 | 0.643 |
| Submissive/Emotional urination (SubEmoUrn) | Urinates when approached, petted, handled or picked up | 397 | 0.202 | 0.594 |
| Separation-related urination (SepUrn) | Urinates when left alone at night, or during the daytime | 397 | 0.385 | 0.832 |
| Separation-related defecation (SepDef) | Defecates when left alone at night, or during the daytime | 397 | 0.317 | 0.752 |
| Hyperactivity (Hyper) | Hyperactive, restless, has trouble settling down | 397 | 0.897 | 1.179 |
| Staring | Stares intently at nothing visible | 393 | 0.583 | 0.981 |
| Fly-snapping (SnapFlies) | Snaps at (invisible) flies | 390 | 0.274 | 0.716 |
| Tail-chasing (TailChase) | Chases own tail/hind end | 392 | 0.367 | 0.785 |
| Light/Shadow-chasing (ShadowChase) | Chases/follows shadows, light spots, etc. | 382 | 0.539 | 1.071 |
| Barking | Barks persistently when alarmed or excited | 397 | 1.879 | 1.482 |
| Grooming self (GroomSelf) | Licks him/herself excessively | 397 | 1.128 | 1.268 |
| Grooming others (GroomOthers) | Licks people or objects excessively | 397 | 0.937 | 1.241 |
| Other stereotypic behavior (OtherBehaviors) | Displays other bizarre, strange, or repetitive behavior(s) | 361 | 0.773 | 1.331 |

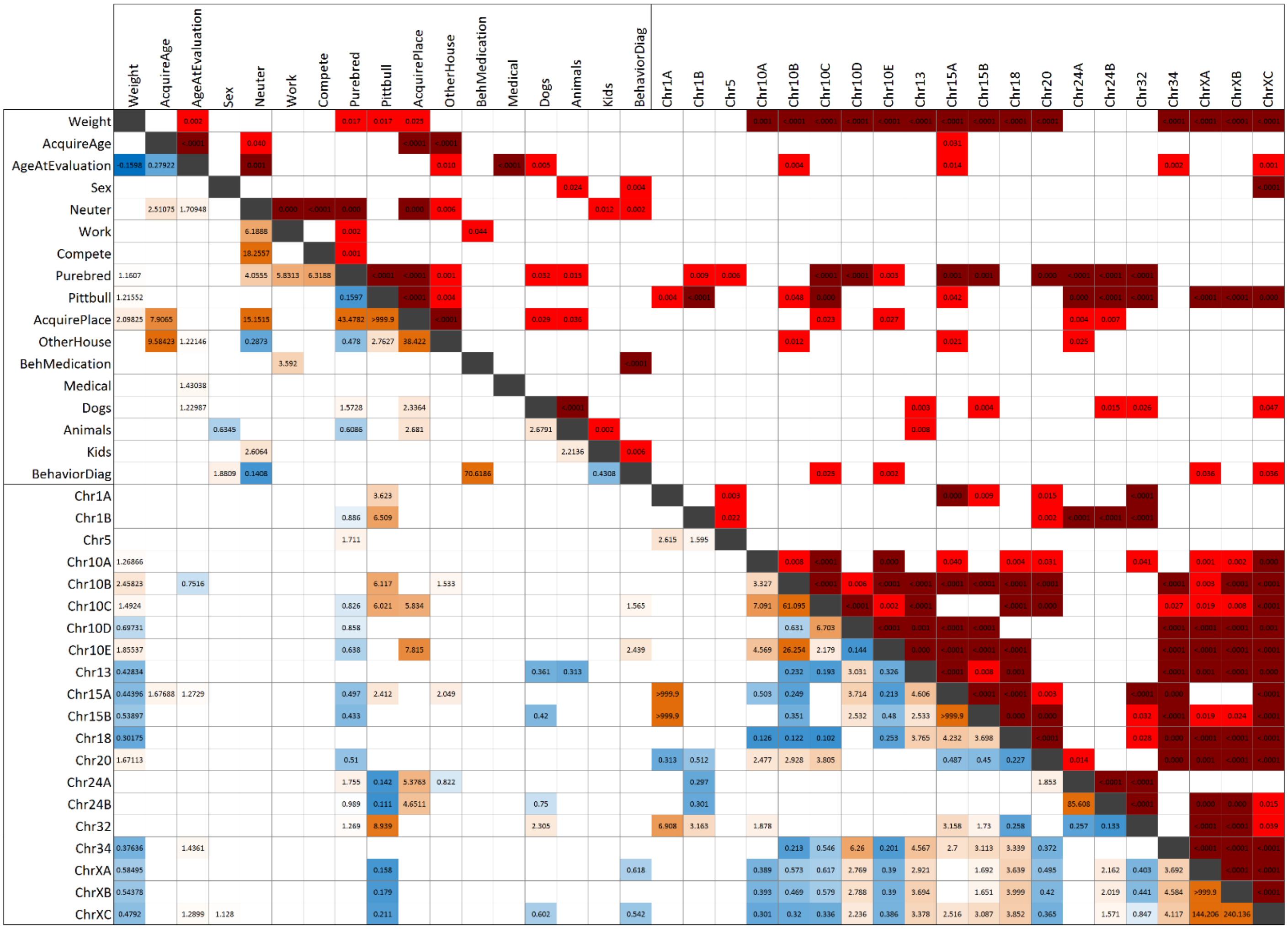


**Supplementary Figure S1.** Duplicate of Figure 1, but with p-values (which require too small of a font size to include in main article). Pairwise association of questionnaire variables and genetic markers. Significance test is shown above the diagonal line and effect size and direction below (odds ratio for categorical variables and estimate ratio for continuous variables). SNP alleles are given according to the CanFam3 nomenclature: Reference allele is A and Alternative is B. Genetic marker significance test and correlation are for AA vs BB. In the top right, red denotes significant association, p≤0.05; and dark red is significant association, p≤0.001. In the bottom left, red is positive association and blue negative. Values are colored in a gradient from red to blue according to their value. Only values with a significant association are displayed.

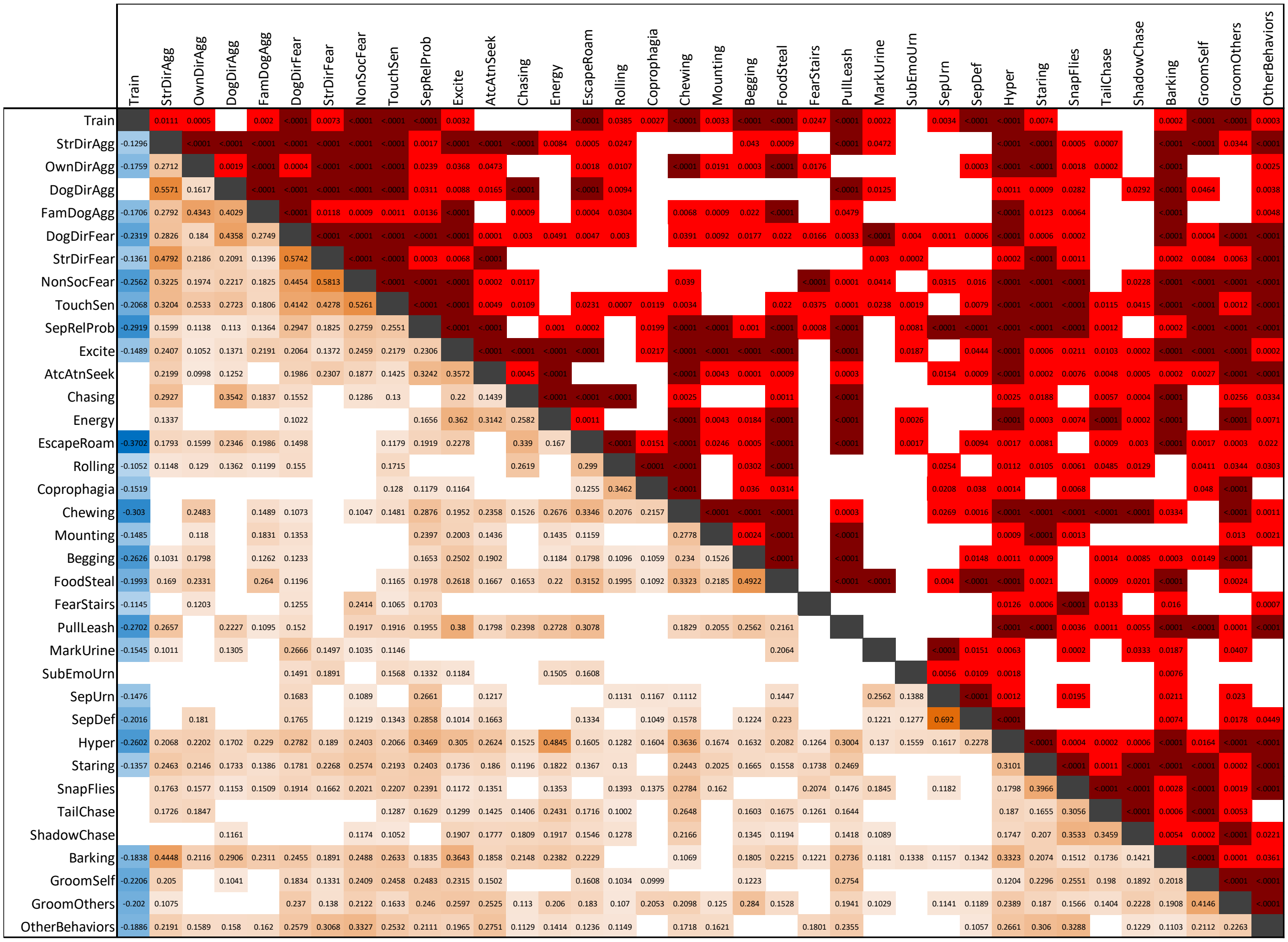


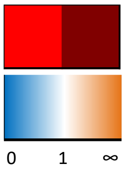


**Supplementary Figure S2.** Correlation table of C-BARQ variables. Significance test (Top right) Pearson's correlation coefficients and direction (Bottom left). The correlation among problematic behaviors measured by the C-BARQ tool is very evident. For the most part dogs displaying problematic behavior tend to show more than one distinct type of it. The variable Trainability is the only positive trait among all of them and is the only one that consistently displays the opposite direction. In the top right, red denotes significant association, p≤0.05; and dark red is significant association, p≤0.001. In the bottom left, red is positive association and blue negative. Values are colored in a gradient from red to blue according to their value. Only values with a significant association are displayed.

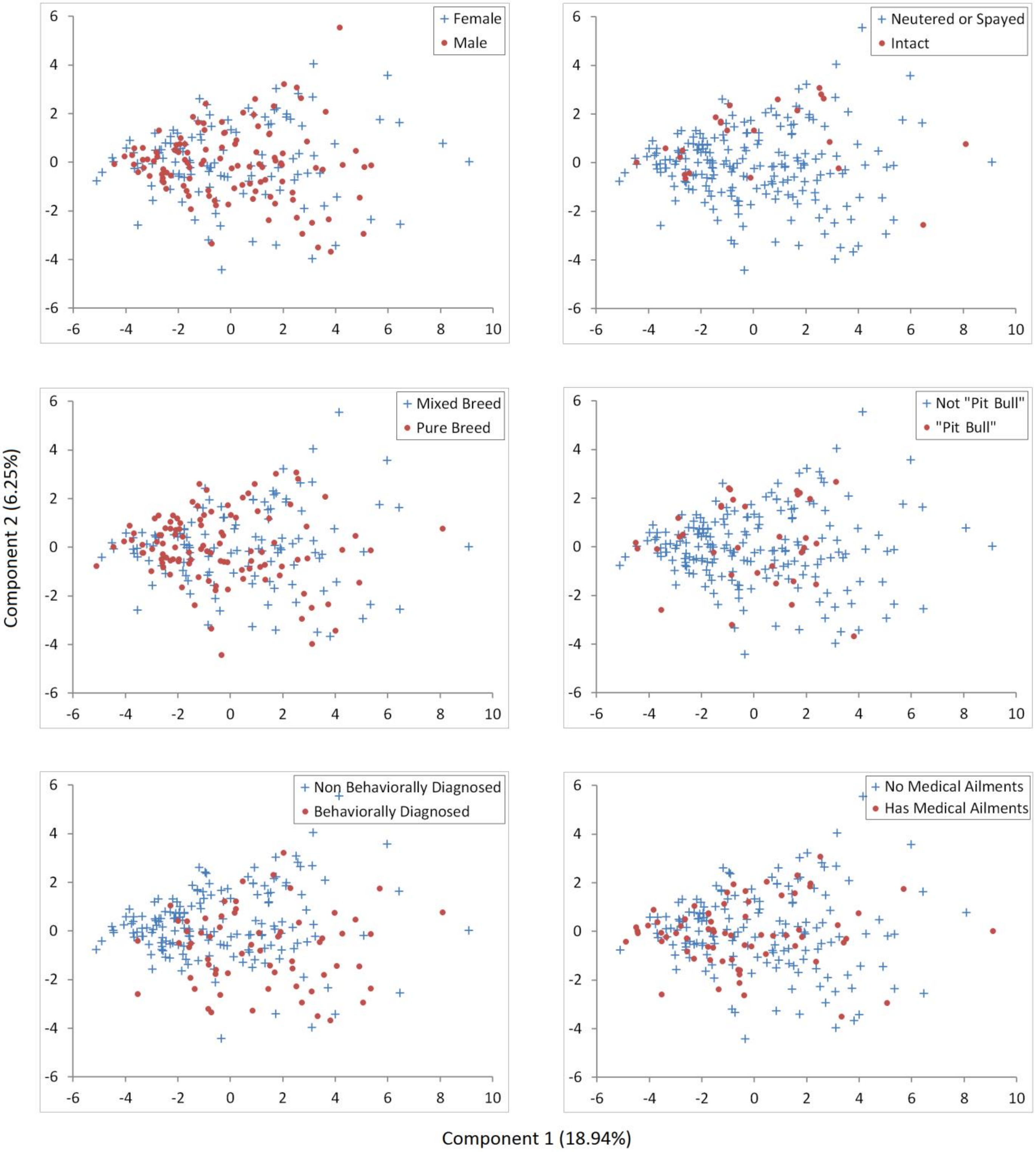


**Supplementary Figure S3.** Principal components analysis observation scores of C-BARQ behavioral traits. Observations in each panel are represented differently to represent the observation distribution for each categorical variable displayed. This allows us to detect effects that are suggestive of variable associations or sampling bias. Only the first two components are shown.

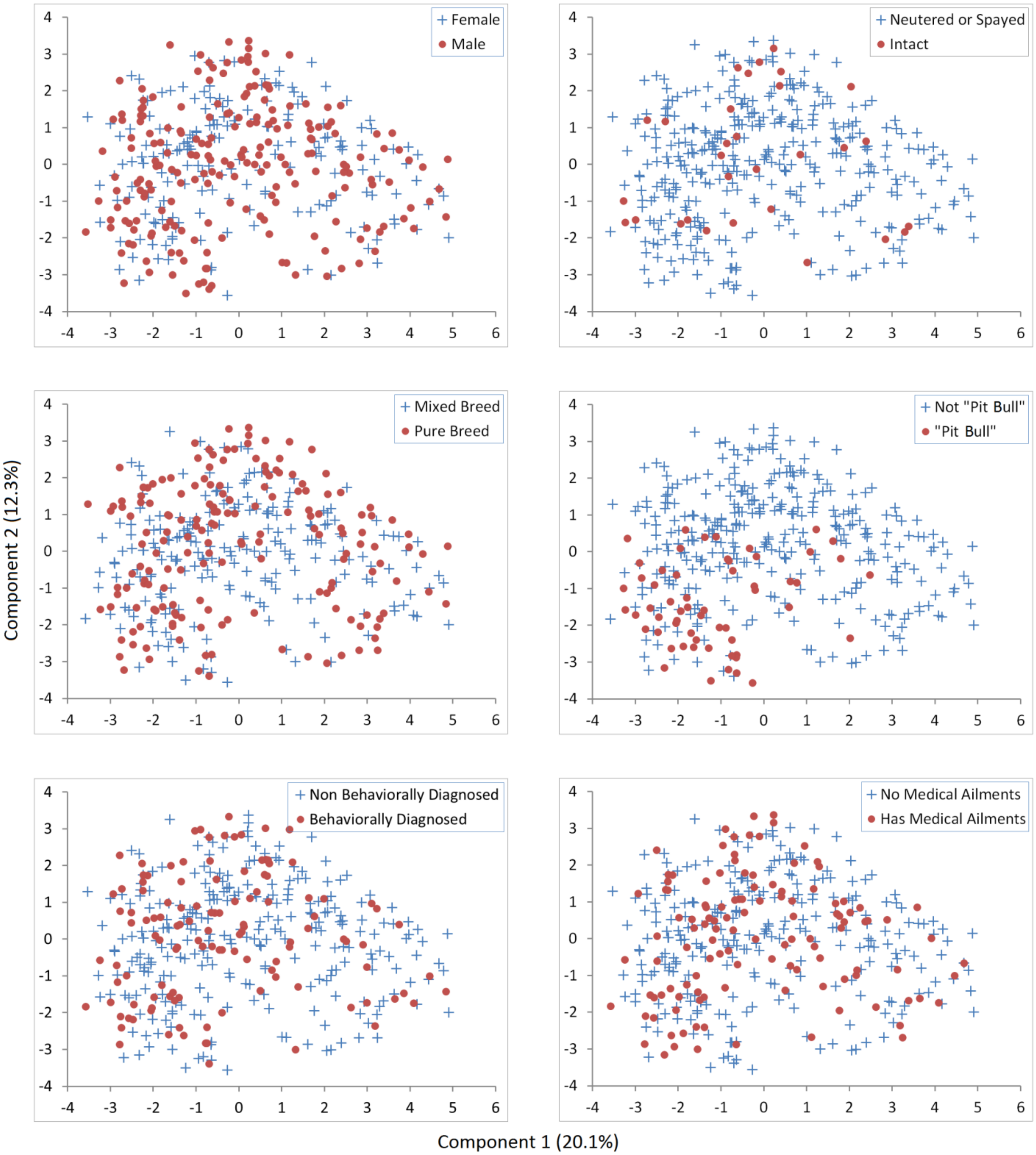


**Supplementary Figure S4.** Principal components analysis observation scores of genetic markers. Observations in each panel are represented differently to represent the observation distribution for each categorical variable displayed. This allows us to detect effects that are suggestive of variable associations or sampling bias. Only the first two components are shown.

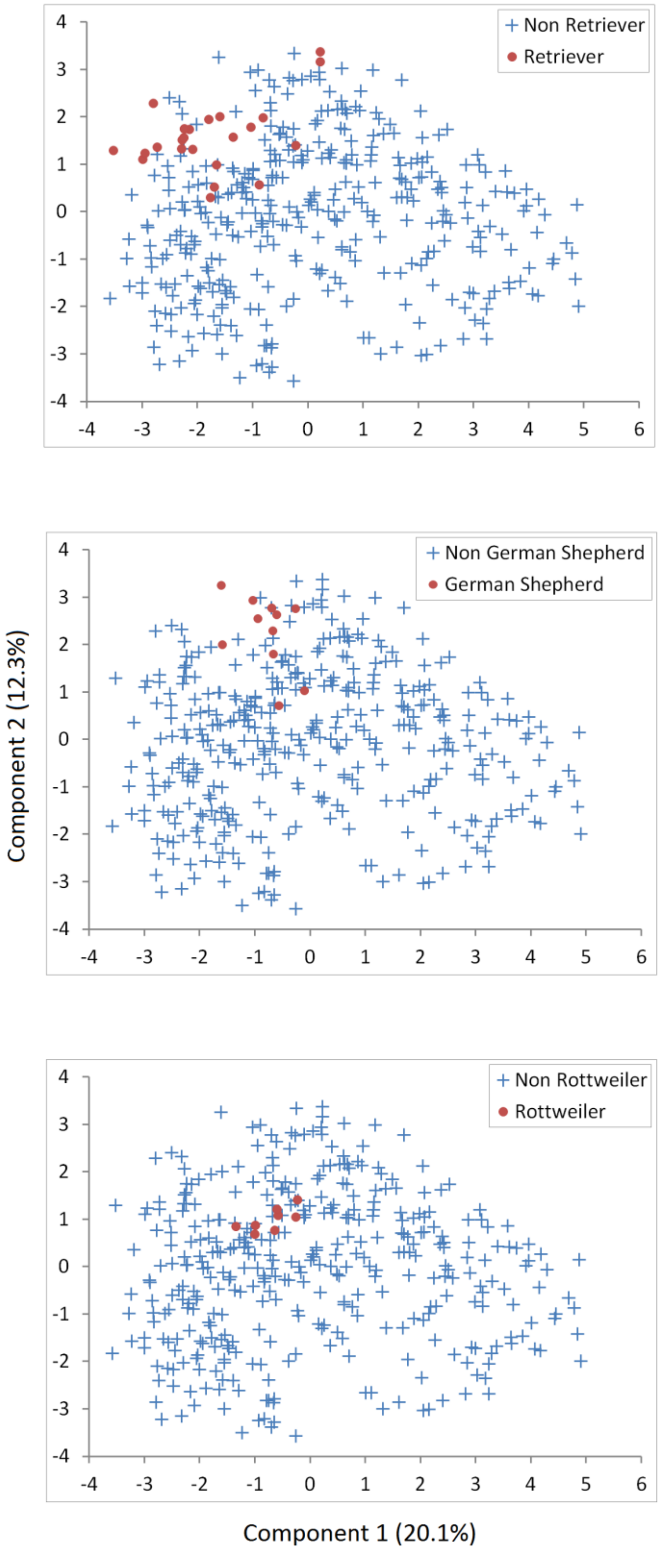


**Supplementary Figure S5.** Principal components analysis observation scores of genetic markers. Additional breed pattern examples. Observations in each panel are represented differently to represent the observation distribution for each categorical variable displayed. These panels are an extension of what was observed in the previous plots where Pitbull designation did show a bias while mixed breeds did not. To further evaluate that effect we have these three additional plots: Retrievers (Labrador Retrievers, Golden Retrievers and Curly Coat Retrievers), German Shepherds and Rottweilers. Only the first two components are shown.

| | | Chr1A | Chr1B | Chr5 | Chr10A | Chr10B | Chr10C | Chr10D | Chr10E | Chr13 | Chr15A | Chr15B | Chr18 | Chr20 | Chr24A | Chr24B | Chr32 | Chr34 | ChrXA | ChrXB | ChrXC |
|---|---|---|---|---|---|---|---|---|---|---|---|---|---|---|---|---|---|---|---|---|---|
| | Behavioral Diagnosed | | | | | 0.016 | | | 0.004 | 0.046 | | | | | | | | | 0.030 | 0.022 | |
| | Behavioral Diagnosed Pitbull | | | | | | | | | | | | | | | | | | | | |
| | Behavioral Medication | | | | | | | | | | 0.019 | | | | | | | | | | |
| By Diagnosis | Anxiety | 0.025 | | | | | | | | | | | | 0.022 | 0.031 | | | | | | |
| | Fear | | | | | | | | | | | | | | | | | | | | |
| | Aggression | | | | | | | | 0.017 | | | | | | | | | | | | |
| | Compulsive | | | | | | | | | | | | | | | | | | | | |
| | Sleep | | | | | | | | | | | | | | | | | | | | |

**Supplementary Figure S6.** Duplicate of Figure 3, but with p-values (which require too small of a font size to include in main article). Diagnostic prediction. Top shows significant marker prediction of a behavioral diagnosis and medication usage. Bottom shows significant marker prediction of specific behavioral diagnoses.

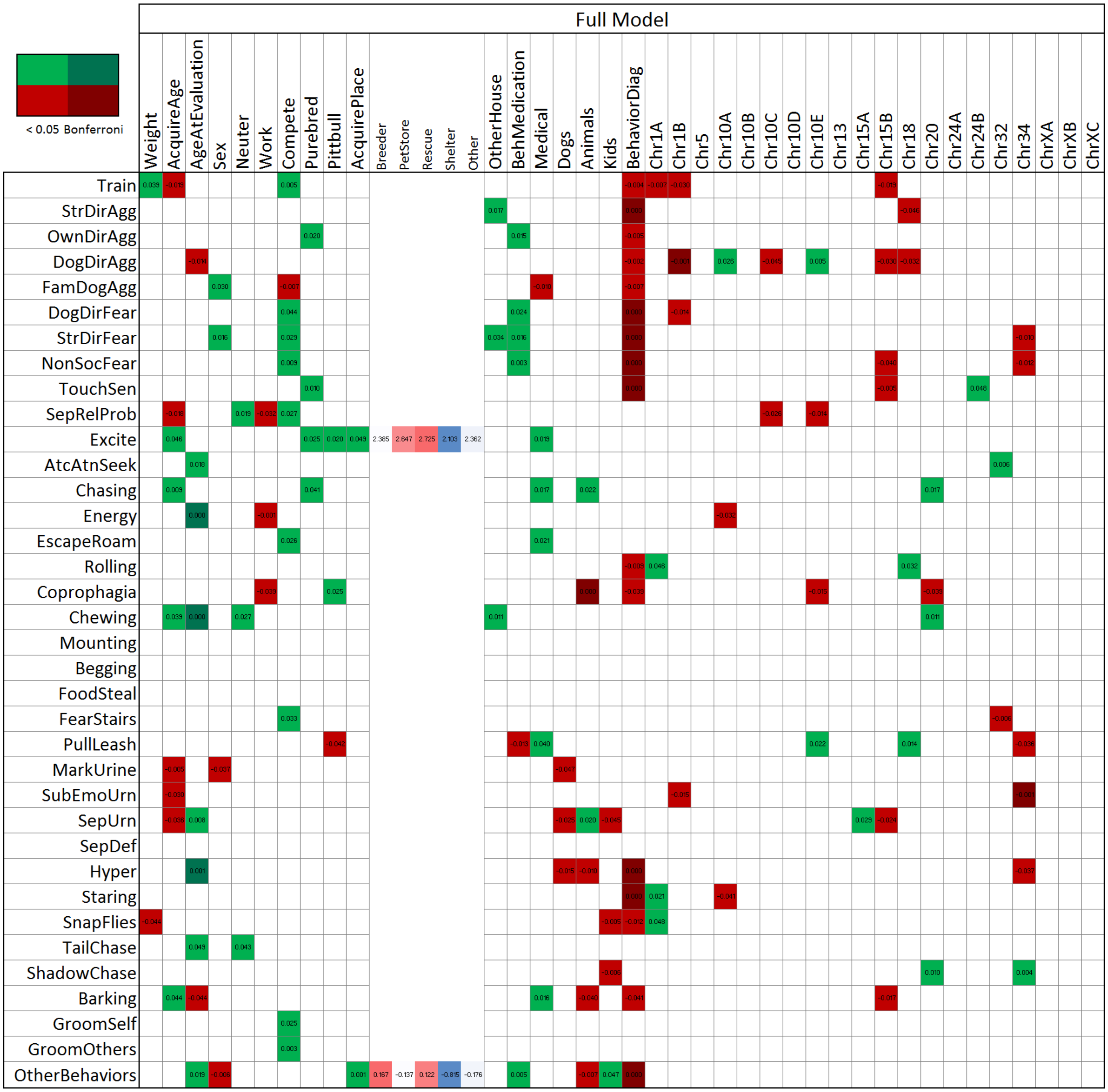


**Supplementary Figure S7.** Duplicate of Figure 4, but with p-values (which require too small of a font size to include in main article). Full Model Mode (FMM): generalized linear model associations of C-BARQ behavioral traits by questionnaire and genetic markers evaluated together. Each behavioral trait was modeled but only significant effects are highlighted. SNP alleles are given according to the CanFam3 nomenclature: Reference allele is A and Alternative is B. Green denotes decreased risk and red increased risk of the A vs. the B allele. A darker shade of green or red denotes significant at a Bonferroni level adjusted by trait. When the effect of place acquired (AcquirePlace) is significant, the Least Square Mean estimate of each of its levels is shown in the columns to its right; color gradient is arranged from lowest to largest.

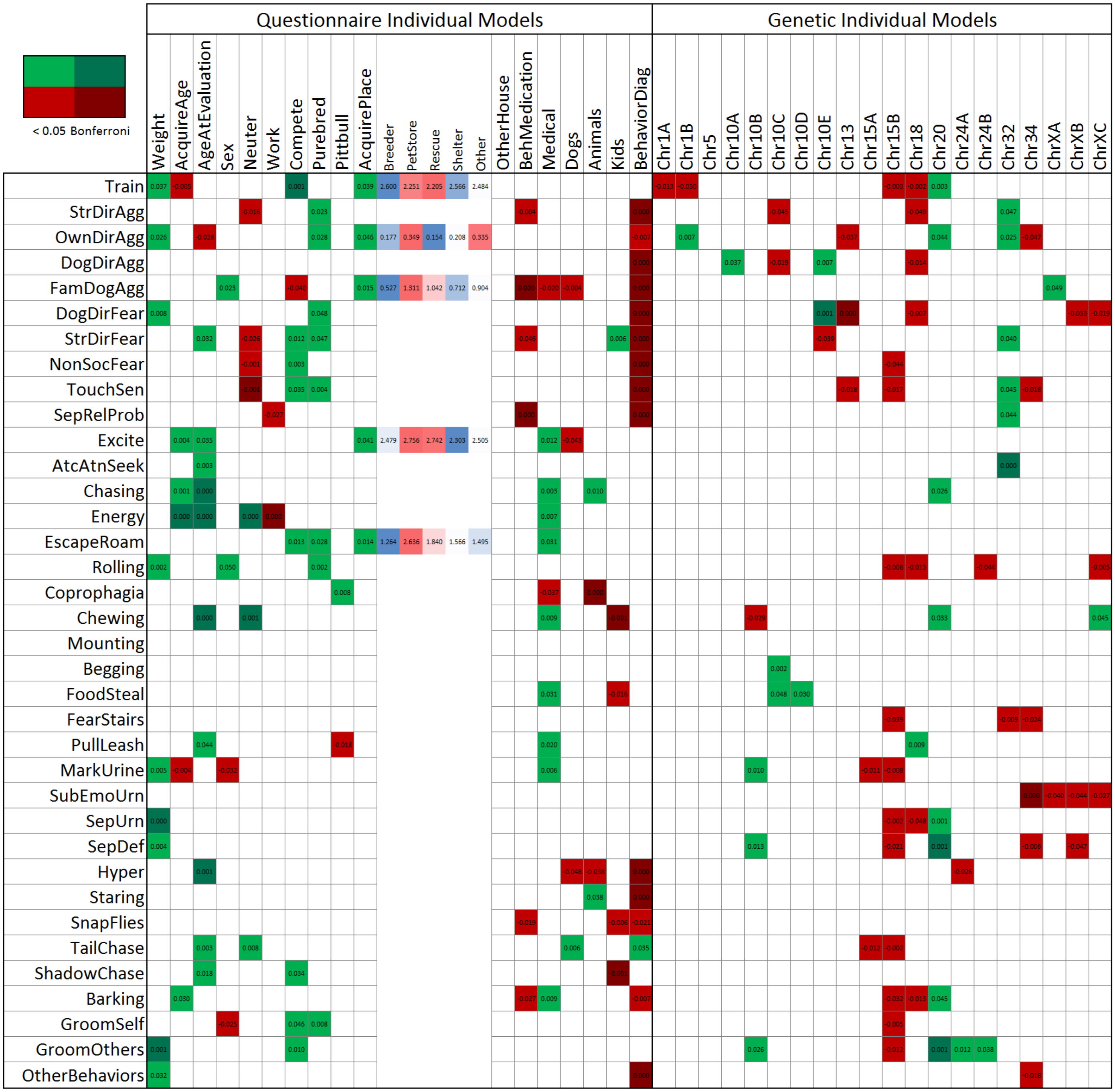


**Supplementary Figure S8.** Duplicate of Figure 5, but with p-values (which require too small of a font size to include in main article). Individual Model Mode (IMM): generalized linear model associations of C-BARQ behavioral traits by questionnaire and genetic markers evaluated individually. Each behavioral trait was modeled but only significant effects are highlighted. SNP alleles are given according to the CanFam3 nomenclature: Reference allele is A and Alternative is B. Green denotes decreased risk and red increased risk of the A vs. the B allele. Green denotes decreased risk and red increased risk. A darker shade of green or red denotes significant at a Bonferroni level adjusted by trait. When the effect of acquired place (AcquirePlace) is significant, the Least Square Mean estimate of each of its levels is shown in the columns to its right; color gradient is arranged from lowest to largest.

## 50%

| | Weight (Kg) | AcquirAGE | AgeAtEvaluation | Sex (Female vs. Male) | Neuter (Fixed vs. Intact) | Work (No vs. Yes) | Compete (No vs. Yes) | Purebred (No vs. Yes) | Pittbull (No vs. Yes) | AcquirPLACE (Breeder vs. Shelter) | AcquirPLACE (Other vs. Shelter) | AcquirPLACE (PetStore vs. Rescue) | AcquirPLACE (Rescue vs. Shelter) | OtherHOUSE (No vs. Yes) | Medication (No vs. Yes) | Medical (No vs. Yes) | Dogs (No vs. Yes) | Animals (No vs. Yes) | Kids (No vs. Yes) | Behavior (No vs. Yes) | Chr1A | Chr1B | Chr5 | Chr10A | Chr10B | Chr10C | Chr10D | Chr10E | Chr13 | Chr15A | Chr15B | Chr18 | Chr20 | Chr24A | Chr24B | Chr32 | Chr34 | ChrXA | ChrXB | ChrXC | R2 | Rescaled R2 | Concondant | Discordant | Tied |
|---|---|---|---|---|---|---|---|---|---|---|---|---|---|---|---|---|---|---|---|---|---|---|---|---|---|---|---|---|---|---|---|---|---|---|---|---|---|---|---|---|---|---|---|---|---|
| Train | 1.017 | | | | | | 0.171 | | | | | | | | | | | | | | 3.195 | | | | | | | | | | | 1.570 | | 0.683 | | | | 0.503 | | | 0.1038 | 0.1384 | 67.6 | 32.3 | 0.1 |
| StrDirAgg | | 1.002 | | | | | | 0.522 | | | | | | | | | | 0.579 | | 3.290 | | | | | | | | 0.486 | | | | | | | | | | | | | 0.1167 | 0.1556 | 70 | 29.7 | 0.3 |
| OwnDirAgg | | | | | | 0.332 | | | | | | | | | 0.335 | | | | | 2.031 | | | | | | | | | | | | | 0.765 | | | | | | | | 0.0543 | 0.0726 | 52.1 | 27.2 | 20.7 |
| DogDirAgg | | | | | | | 3.690 | | | | | | | | | 0.649 | | | | 2.985 | | 1.944 | | 0.166 | | | | 0.678 | | | | | | | | | | | | | 0.1145 | 0.1528 | 68.1 | 29.7 | 2.2 |
| FamDogAgg | | | | 0.329 | | | | | | | | | | | | | 2.280 | | | 2.904 | | | | | | | | | | 2.890 | | | | | | | | | | | 0.1407 | 0.1879 | 68.7 | 25.9 | 5.4 |
| DogDirFear | 1.021 | | | | | | 0.284 | | | | | | | | | | | | | 3.080 | | 1.978 | | | | | | | | | | | | 1.375 | | | | | | | 0.1031 | 0.1377 | 69.2 | 30.6 | 0.1 |
| StrDirFear | | | | 0.638 | | | 0.124 | | | | | | | | 0.412 | | | 0.588 | | 4.138 | | | | | | | | 0.731 | | | 2.668 | | | | | 0.456 | | | | | 0.1572 | 0.21 | 72.4 | 27 | 0.6 |
| NonSocFear | | | | | | | 0.043 | | | | | | | | | 0.582 | | | | 4.682 | | | | | | | | | | | | | | | 0.786 | | | | | 2.092 | 0.1603 | 0.2142 | 72 | 24.2 | 3.8 |
| TouchSen | | | | | 0.274 | | | | | | | | | | | | | | | 2.798 | | | | | | | | | 1.315 | | | | | | | | | | | | 0.105 | 0.14 | 49.1 | 16.1 | 34.8 |
| SepRelProb | | | | | | | | | | | | | | | 3.787 | | | | | | | 1.840 | | | | 0.645 | | | | | | | | | | | | | | | 0.0542 | 0.0723 | 56.9 | 33.5 | 9.5 |
| Excite | | 1.003 | | | | | | | | | | | | | | | 1.586 | | | | | | 0.476 | | 0.489 | | | | | | | | | | | | | | | | 0.0476 | 0.0636 | 61.4 | 37.9 | 0.8 |
| AtcAtnSeek | | | 1.002 | | | | | | | | | | | | | | | | | 1.497 | | | | | | | | 0.456 | | | | | 0.503 | | | 0.511 | | | | | 0.0826 | 0.1102 | 65.2 | 34.6 | 0.2 |
| Chasing | | 1.003 | | | | | | | | | | | | | | 0.433 | | 0.636 | | | 4.299 | 1.216 | | | | | | | | | | | | | | | | | | | 0.0969 | 0.1292 | 67.3 | 32 | 0.7 |
| Energy | | | 1.004 | 1.573 | | 3.698 | | | 0.566 | | | | | | | | 1.562 | | | | | | | | | | | | | | | | | | | | | | | | 0.1463 | 0.1982 | 72.9 | 26.3 | 0.9 |
| EscapeRoam | | | | | | | 0.252 | | | | | | | | | 0.552 | | | | | | | 0.669 | | | | 0.428 | | | | 2.165 | | | | | | | | | | 0.0773 | 0.1036 | 64.1 | 32.9 | 3 |
| Rolling | 1.019 | | 0.999 | | | | | 0.538 | | | | | | | | | | | | | | | | | | | | | | | | | | | | | | | | | 0.0543 | 0.0734 | 62 | 38 | 0 |
| Coprophagia | | | | 0.614 | | | | | 0.494 | | | | | | | | | 1.765 | | 1.601 | 0.373 | | | | | | | | | | | | | | | | | | | | 0.0639 | 0.0855 | 60.1 | 30.5 | 9.4 |
| Chewing | | 0.996 | 1.005 | | | | | | | | | | | 0.743 | | | | | 2.313 | | | | | | | | | | | | | | 0.786 | | | | | | | 0.587 | 0.1589 | 0.229 | 75.5 | 24.5 | 0 |
| Mounting | | | | | | | | | | | | | | | | | | | | | | | | | | | | | | | | | | | | | | | | | - | - | - | - | - |
| Begging | | | | | | 0.270 | | | | | | | | 0.775 | | | | | | | | 1.847 | | | | 0.477 | | | | | | | | | | | | | | | 0.07 | 0.0986 | 63.9 | 32.1 | 4 |
| FoodSteal | | | | | | | | | | | | | | | | 0.597 | | | | | | | | | | 0.471 | | | | | | | | | | | | | | | 0.0419 | 0.0592 | 50.1 | 30.2 | 19.7 |
| FearStairs | | | | | | | | | | | | | | | | | | | | | | | | | | | | | | | | | | | | 2.620 | | | | | 0.0228 | 0.0344 | 38.4 | 23.2 | 38.5 |
| PullLeash | | 1.002 | | | | | | | 2.286 | | | | | | | 0.631 | | 0.661 | | | | | | | | | | 0.535 | | | | 0.560 | | | | | | | | | 0.0792 | 0.1059 | 66.5 | 33.1 | 0.4 |
| MarkUrine | 1.053 | 0.996 | | 2.908 | | | | | | | | | | | | 0.232 | 1.827 | | | | | | | | | | | | | | | | | | | | | | 0.429 | | 0.1007 | 0.1853 | 76.1 | 23.9 | 0 |
| SubEmoUrn | | | | | | | | | | | | | | | | | | | | | | | | | | | | | | | | 0.579 | | | | | | | | | 0.0211 | 0.039 | 32 | 13.1 | 54.9 |
| SepUrn | 1.049 | | | | | | | | | | | | | | | | | | | | | 2.413 | | | | 1.372 | | | | | 0.728 | | | | | | | | | | 0.0948 | 0.1453 | 70.7 | 29.2 | 0.1 |
| SepDef | | | | | | | | | | | | | | | | | | | | | | | | | 0.538 | | | | | | | | | | | | 1.624 | | | | 0.0405 | 0.0658 | 53.8 | 25.6 | 20.6 |
| Hyper | | | 1.001 | | | | | | | | | | | | | 0.611 | 1.704 | 1.634 | | 3.420 | | | | | | | | | | | | | | 1.894 | | | | | | | 0.1189 | 0.159 | 70.1 | 29.7 | 0.1 |
| Staring | | | | | | | | | | | | | | | | | | 0.655 | | 1.857 | 0.323 | | | | | | | | | | | | | | | | | | | | 0.0408 | 0.057 | 49.9 | 25.9 | 24.1 |
| SnapFlies | | | | | | | | | | | | | | | | | | | | | | | | | | | | | | | | | | | | | | | | | - | - | - | - | - |
| TailChase | 0.976 | | 1.002 | | | | | | | | | | | | | | 0.579 | | | 0.468 | | | | | | | | | | 0.801 | 10.69 | | | | | | | | | | 0.1149 | 0.1743 | 73.3 | 26.7 | 0 |
| ShadowChase | | | | | | | | | | | | | | | | | | | 2.281 | | | | | | | | | | | | | | | | | | | | | | 0.0256 | 0.0376 | 27.2 | 11.8 | 61.1 |
| Barking | | 1.003 | 0.999 | | | | | | | | | | | | | 0.427 | 1.767 | | | 2.427 | | | 0.458 | | | | | | | | 2.144 | | | | | | | | | 1.967 | 0.1399 | 0.1904 | 72.4 | 27.6 | 0 |
| GroomSelf | | | 0.998 | 1.658 | | | | | | | | | | | | | | | | 1.647 | | | | | | | | | | | 2.742 | | | | | | | | | | 0.0721 | 0.1003 | 65.8 | 33.7 | 0.5 |
| GroomOthers | | | | | | | 0.184 | | | | | | | | | | | | | | | | | | | 0.554 | | | | | | | 0.989 | | | | | | | | 0.0605 | 0.081 | 57 | 30.2 | 12.8 |
| OtherBehaviors | | | 1.001 | | | | | | | | | | | | | | | | | 3.342 | | | | | | | | | | | 2.613 | | | | | | 3.477 | | | | 0.107 | 0.1518 | 69.1 | 30.3 | 0.6 |

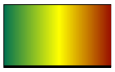


**Supplementary Figure S9.** Logistic regression model selection using a stepwise selection method for traits recoded as case/control: case threshold at 50th percentile of the data. Odds ratio values are used to determine effect size and direction. Predicted probabilities and observed responses are displayed for each row. SNP alleles are given according to the CanFam3 nomenclature: Reference allele is A and Alternative is B. Green denotes decreased risk and red increased risk of the A vs. the B allele.

## 75%

| | Weight | AcquirAGE | AgeAtEvaluation | Sex (Female vs. Male) | Neuter (Fixed vs. Intact) | Work (No vs. Yes) | Compete (No vs. Yes) | Purebred (No vs. Yes) | Pittbull (No vs. Yes) | AcquirPLACE (Breeder vs. Shelter) | AcquirPLACE (Other vs. Shelter) | AcquirPLACE (PetStore vs. Rescue) | AcquirPLACE (Rescue vs. Shelter) | OtherHOUSE (No vs. Yes) | Medication (No vs. Yes) | Medical (No vs. Yes) | Dogs (No vs. Yes) | Animals (No vs. Yes) | Kids (No vs. Yes) | Behavior (No vs. Yes) |
|---|---|---|---|---|---|---|---|---|---|---|---|---|---|---|---|---|---|---|---|---|
| Train | | | | | | | 0.230 | | | | | | | | | | | | | |
| StrDirAgg | | 1.003 | | | | | | | | | | | | | | | | | | 5.416 |
| OwnDirAgg | 1.044 | | 0.998 | | | | | | 0.317 | | | | | | | | | | | |
| DogDirAgg | | | | | | | | | | | | | | | | | | | | 3.432 |
| FamDogAgg | | | | 0.384 | | | 3.746 | | | | | | | | | | 3.289 | | | 4.990 |
| DogDirFear | | | | | | | | | | | | | | | | | | | | 4.273 |
| StrDirFear | | | | | | | | | | | | | | | | | | | | 3.866 |
| NonSocFear | | | | | | | | | | | | | | | | | | | | 5.584 |
| TouchSen | | | | | | | | | | | | | | | | | | | | 3.373 |
| SepRelProb | | | | | | | | | | | | | | | 3.424 | | 0.543 | | | 1.783 |
| Excite | | | | | | 3.735 | | | 0.487 | 1.726 | 1.025 | 0.376 | 0.273 | 0.342 | | | 1.806 | | | 1.746 |
| AtcAtnSeek | | | 1.002 | | | | | | | | | | | | | | | | | |
| Chasing | | | | | | | 2.748 | | | | | | | | | | | 0.532 | | 1.770 |
| Energy | | | 1.004 | | | 5.700 | | | | | | | | | | | | | | |
| EscapeRoam | | | | | | | | | | 3.537 | 1.407 | 0.221 | 1.289 | | | | | | | |
| Rolling | | | | 0.579 | | | | | | | | | | | | | | | | |
| Coprophagia | | | | | | 3.198 | | | 0.144 | | | | | 0.623 | | 1.786 | 2.154 | 2.707 | | 1.817 |
| Chewing | | | 1.003 | | | | | | | | | | | | | | | | | |
| Mounting | | | | | | | | | | | | | | | | | | | | |
| Begging | | | | | | | | | | | | | | | | | | | | |
| FoodSteal | | | | | | | | | | | | | | | | | | | | |
| FearStairs | | | | | | | | | | | | | | | | | | | | |
| PullLeash | | | 1.002 | | | | | | 2.116 | | | | | | | | | | | |
| MarkUrine | 1.053 | 0.996 | | 2.908 | | | | | | | | | | | | 0.232 | 1.827 | | | |
| SubEmoUrn | | | | | | | | | | | | | | | | | | | | |
| SepUrn | 1.049 | | | | | | | | | | | | | | | | | | | |
| SepDef | | | | | | | | | | | | | | | | | | | | |
| Hyper | | | 1.004 | | | | | | | | | | | | | | | | | |
| Staring | | | | | | | | | | | | | | | | | | 0.453 | | 2.893 |
| SnapFlies | | | | | | | | | | | | | | | | | | | | |
| TailChase | 0.976 | | 1.002 | | | | | | | | | | | | | | 0.579 | | | 0.468 |
| ShadowChase | | | 1.002 | | | | | | | | | | | | | | | | 3.207 | |
| Barking | | | | | | | | | | | | | | | | | | 1.706 | | 2.515 |
| GroomSelf | | | | | | | | | | | | | | | | | | | | |
| GroomOthers | 1.049 | | 1.002 | | | | | | | | | | | | | | | | | |
| OtherBehaviors | | | 1.001 | | | | | | | | | | | | | | | | | 3.548 |

| | Chr1A | Chr1B | Chr5 | Chr10A | Chr10B | Chr10C | Chr10D | Chr10E | Chr13 | Chr15A | Chr15B | Chr18 | Chr20 | Chr24A | Chr24B | Chr32 | Chr34 | ChrXA | ChrXB | ChrXC |
|---|---|---|---|---|---|---|---|---|---|---|---|---|---|---|---|---|---|---|---|---|
| Train | 4.098 | 2.639 | 0.576 | 0.708 | | | | | | | 2.000 | | | | | | | | | |
| StrDirAgg | | | | | | 2.021 | | | | | | 3.109 | | 2.302 | | | | | | |
| OwnDirAgg | | | | | | | | | | | | | | | | 0.323 | | | | 0.409 |
| DogDirAgg | | | | | | | | 0.511 | | | | 1.096 | | | | | | | | |
| FamDogAgg | | | | | | | | | | 1.925 | | | | | | | | | | |
| DogDirFear | | | | | | | | 0.398 | | | | 3.171 | | | | | | | | 2.586 |
| StrDirFear | | | | | | | | | | | | | | | | | | | | |
| NonSocFear | 3.594 | | | | | | | | | | | | | | | | 1.817 | | | |
| TouchSen | | | | | | | | | | | 3.429 | | | | | | 2.701 | | | |
| SepRelProb | | | | | | | | | | | | | | | | | | | | |
| Excite | | | | | | | | | | 3.022 | | | | | | | | | | |
| AtcAtnSeek | | | | | | | | 0.603 | | | | | | | | 0.452 | | | | |
| Chasing | | | | | | | | 2.065 | | 1.411 | | | | | | | | | | |
| Energy | | | | | | | | | | | | | | | | | | | | |
| EscapeRoam | | | | | | | | | | | | | | | | | | | | |
| Rolling | | | | | | | | | | | | | | | | | | | | 2.431 |
| Coprophagia | | | | | | | | | | 1.862 | | | | | | | | | | |
| Chewing | | | | | | | | | | | | | 0.391 | | | | | | | 0.396 |
| Mounting | | | | | | | | | | | | | | | | | | | | |
| Begging | 2.841 | | | | | | | | | | | | | | 0.430 | | | | | |
| FoodSteal | | | | | | | | | | | | | | | | | | | | |
| FearStairs | | | | | | | | | | | | | | | | 2.620 | | | | |
| PullLeash | | | | | | | | 0.710 | | | | | | | | | 3.952 | | | |
| MarkUrine | | | | | | | | | | | | | | | | | | | 0.429 | |
| SubEmoUrn | | | | | | | | | | | | 0.579 | | | | | | | | |
| SepUrn | | 2.413 | | | | 1.372 | | | | | 0.728 | | | | | | | | | |
| SepDef | | | | | 0.538 | | | | | | | | | | | | 1.624 | | | |
| Hyper | | | | | | | | | | | | | | | | | | | | |
| Staring | | | | | | | | | | | | | | | | | | | | |
| SnapFlies | | | | | | | | | | | | | | | | | | | | |
| TailChase | | | | | | | | | | 0.801 | 10.69 | | | | | | | | | |
| ShadowChase | | | | | | 0.444 | | | | | | | | | | | 0.426 | | | |
| Barking | | | | | | | | | | | | | | | | | 2.851 | | | 1.705 |
| GroomSelf | | | | | | | | | | | 2.467 | | | | 0.465 | | | | | |
| GroomOthers | | | | | | | | | | | | | 0.587 | 0.316 | | | | | | |
| OtherBehaviors | | | | | | | | | | | 2.900 | | | | | | 4.432 | | | |

| | R2 | Rescaled R2 | Concondant | Discordant | Tied |
|---|---|---|---|---|---|
| Train | 0.098 | 0.1483 | 69.6 | 27.5 | 2.9 |
| StrDirAgg | 0.1633 | 0.244 | 76.2 | 23.5 | 0.3 |
| OwnDirAgg | 0.0885 | 0.1343 | 70.3 | 29.7 | 0 |
| DogDirAgg | 0.101 | 0.1521 | 66.9 | 24.4 | 8.8 |
| FamDogAgg | 0.141 | 0.218 | 73.4 | 22.1 | 4.5 |
| DogDirFear | 0.1288 | 0.211 | 73.9 | 22.8 | 3.2 |
| StrDirFear | 0.0735 | 0.1101 | 41.4 | 10.7 | 47.9 |
| NonSocFear | 0.1342 | 0.2055 | 63.8 | 16.1 | 20.1 |
| TouchSen | 0.0923 | 0.137 | 63.1 | 26.8 | 10.1 |
| SepRelProb | 0.0647 | 0.0962 | 52.7 | 23.2 | 24.1 |
| Excite | 0.106 | 0.1611 | 70.5 | 28.6 | 0.8 |
| AtcAtnSeek | 0.0636 | 0.0978 | 66.6 | 32.7 | 0.7 |
| Chasing | 0.0668 | 0.1069 | 66.6 | 30.7 | 2.7 |
| Energy | 0.0821 | 0.1417 | 69 | 26.3 | 4.7 |
| EscapeRoam | 0.0509 | 0.0961 | 52.2 | 21.4 | 26.4 |
| Rolling | 0.0411 | 0.0624 | 53.8 | 26.8 | 19.4 |
| Coprophagia | 0.1123 | 0.193 | 75.5 | 23.8 | 0.7 |
| Chewing | 0.0638 | 0.1228 | 71 | 28.1 | 1 |
| Mounting | - | - | - | - | - |
| Begging | 0.0331 | 0.0579 | 49.2 | 23.8 | 27.1 |
| FoodSteal | - | - | - | - | - |
| FearStairs | 0.0228 | 0.0344 | 38.4 | 23.2 | 38.5 |
| PullLeash | 0.0648 | 0.1 | 66.7 | 32.7 | 0.6 |
| MarkUrine | 0.1007 | 0.1853 | 76.1 | 23.9 | 0 |
| SubEmoUrn | 0.0211 | 0.039 | 32 | 13.1 | 54.9 |
| SepUrn | 0.0948 | 0.1453 | 70.7 | 29.2 | 0.1 |
| SepDef | 0.0405 | 0.0658 | 53.8 | 25.6 | 20.6 |
| Hyper | 0.0388 | 0.0797 | 66.2 | 28.4 | 5.4 |
| Staring | 0.0587 | 0.0974 | 55.7 | 20.3 | 24 |
| SnapFlies | - | - | - | - | - |
| TailChase | 0.1149 | 0.1743 | 73.3 | 26.7 | 0 |
| ShadowChase | 0.0708 | 0.1283 | 70.2 | 29.2 | 0.7 |
| Barking | 0.0663 | 0.1048 | 65.8 | 28.5 | 5.7 |
| GroomSelf | 0.0323 | 0.0544 | 56.4 | 30.8 | 12.9 |
| GroomOthers | 0.0904 | 0.1616 | 73.7 | 26.3 | 0 |
| OtherBehaviors | 0.104 | 0.1575 | 70.1 | 29.3 | 0.6 |

Protective — Detrimental

**Supplementary Figure S10.** Logistic regression model selection using a stepwise selection method for traits recoded as case/control: case threshold at 75th percentile of the data. Odds ratio values are used to determine effect size and direction. Predicted probabilities and observed responses are displayed for each row.

90%

| | Weight | AcquirAGE | AgeAtEvaluation | Sex (Female vs. Male) | Neuter (Fixed vs. Intact) | Work (No vs. Yes) | Compete (No vs. Yes) | Purebred (No vs. Yes) | Pittbull (No vs. Yes) | AcquirPLACE (Breeder vs. Shelter) | AcquirPLACE (Other vs. Shelter) | AcquirPLACE (PetStore vs. Rescue) | AcquirPLACE (Rescue vs. Shelter) | OtherHOUSE (No vs. Yes) | Medication (No vs. Yes) | Medical (No vs. Yes) | Dogs (No vs. Yes) | Animals (No vs. Yes) | Kids (No vs. Yes) | Behavior (No vs. Yes) | Chr1A | Chr1B | Chr5 | Chr10A | Chr10B | Chr10C | Chr10D | Chr10E | Chr13 | Chr15A | Chr15B | Chr18 | Chr20 | Chr24A | Chr24B | Chr32 | Chr34 | ChrXA | ChrXB | ChrXC | R2 | Rescaled R2 | Concondant | Discordant | Tied |
|---|---|---|---|---|---|---|---|---|---|---|---|---|---|---|---|---|---|---|---|---|---|---|---|---|---|---|---|---|---|---|---|---|---|---|---|---|---|---|---|---|---|---|---|---|---|
| Train | | | | | | | 0.230 | | | | | | | | | 0.486 | | | | | | | 0.627 | | | | | | | | | | 0.495 | | | 1.686 | | | | | 0.0956 | 0.2116 | 78.5 | 17.8 | 3.7 |
| StrDirAgg | | | | | | | | | | | | | | | | | | | | 5.844 | | | | | | | | | | | | 1.162 | | | | | | | 0.360 | | 0.107 | 0.2255 | 75 | 15.6 | 9.4 |
| OwnDirAgg | 1.037 | | | | | | | 0.364 | | | | | | | | | | | | | | 0.209 | | | | | | | 4.163 | | | | | | | | | | 0.302 | | 0.085 | 0.1838 | 77.2 | 22.7 | 0.1 |
| DogDirAgg | | | | | | | | | | | | | | | | | | | | 2.607 | | | | | | | | | | | | | | | | | | | | | 0.0196 | 0.0409 | 36.1 | 13.9 | 50 |
| FamDogAgg | | | | | | | 8.876 | | | 1.956 | 0.382 | 0.022 | 0.089 | | | 3.862 | | 5.881 | | 7.443 | | | | | | | | 1.552 | | | | | | | | | | | | | 0.1456 | 0.3905 | 87.7 | 11.8 | 0.5 |
| DogDirFear | | | | 2.219 | | | | | | | | | | | | 0.410 | | | | 4.645 | | | | | | | | 0.209 | | | | | | | | | | | | 1.735 | 0.0881 | 0.1844 | 75.5 | 22.6 | 1.9 |
| StrDirFear | | | | | | | | | | | | | | | | | | | | 3.948 | | | | | | | | | | | | | | | | | | | | | 0.0372 | 0.081 | 43.5 | 11 | 45.5 |
| NonSocFear | | | | | | | | | | | | | | | | | | | | 6.951 | | | | | | | | | | | 4.372 | 3.577 | | | | | 1.397 | | | | 0.1116 | 0.2332 | 77.1 | 18 | 4.8 |
| TouchSen | | | | | | | | | 0.139 | | | | | | | | | | | 3.012 | 3.835 | | | | | | | | | | | | | | | | 5.136 | | | | 0.067 | 0.1524 | 67.2 | 17.2 | 15.6 |
| SepRelProb | | 0.997 | | | | | | | | | | | | | | | | | | 3.280 | | | | | | 3.009 | | | | | | | | | | | | | | | 0.0609 | 0.1293 | 72.7 | 26.6 | 0.7 |
| Excite | | | | | | | | | | | | | | 0.521 | | | | | | | | | | | 0.247 | | | | | | | 0.287 | | | | | | | | | 0.049 | 0.1309 | 68.7 | 22.3 | 9 |
| AtcAtnSeek | | | | | 3.938 | | | | | | | | | | 2.822 | | | | | | | | | | | | | | | | | | | | | 0.247 | | | | | 0.0385 | 0.0929 | 58.2 | 17.4 | 24.4 |
| Chasing | | | | | | | | | | | | | | | | | | | | 2.313 | | | | | | | | | | | | | | | 0.233 | | | | | | 0.0494 | 0.1161 | 66 | 19 | 15 |
| Energy | | | | | | 8.821 | | | | | | | | | | | | | | | | | | | | | | | | | | | | | | | | | | | 0.0308 | 0.0708 | 18.3 | 2.1 | 79.7 |
| EscapeRoam | | | | | | | | | | | | | | | | | | | | | | | | | | | | | | | | | | | | | | | | | - | - | - | - | - |
| Rolling | | | | | | | | | | | | | | | | | | | | | | | | | | | | | | | | | | | | | | | | | - | - | - | - | - |
| Coprophagia | | | | | | | | | | | | | | | | | | 2.529 | | | | | | | | | | | | 0.773 | | | | | | | | | | | 0.0252 | 0.0628 | 58.1 | 25.9 | 16 |
| Chewing | | | | | 4.823 | | | | | | | | | | | | | | | | | | | | | | | | | | | | | | | | | | | | 0.0196 | 0.0514 | 22.7 | 4.7 | 72.6 |
| Mounting | | | | | | | | | | | | | | | | | | | | | | | | | | | | | | | | | | | | | | | | | - | - | - | - | - |
| Begging | | | | | | | | | | | | | | | | | | | | | | | | | | | | | | | | | | | | | | | | | - | - | - | - | - |
| FoodSteal | | | | | | | | | | | | | | | | | | | | | | | | | | | | | | 0.796 | | | | | | | | | | | 0.0186 | 0.0447 | 47.1 | 20.6 | 32.3 |
| FearStairs | | | | | | | | | | | | | | | | | 0.216 | | | | | | | | | | | | | | | 4.193 | | | | 4.562 | 8.546 | | | | 0.0643 | 0.1953 | 75.8 | 19.8 | 4.4 |
| PullLeash | | | | | | | | | | | | | | | | | | | | | | | | | | | | | | | | | | | | | | | | | - | - | - | - | - |
| MarkUrine | | 0.995 | | | | | | | | | | | | | | | | | | | | | | | | | | | | | | | | | | | | | | | 0.0264 | 0.069 | 59.2 | 35.4 | 5.4 |
| SubEmoUrn | | | | | | | | | | | | | | | | | | | | | | | | | | | | | | | | | | | | | | | | | - | - | - | - | - |
| SepUrn | | | | | | | | | | | | | | | | | | | | | | | | | | | | | | | | | | | | | | | | | - | - | - | - | - |
| SepDef | | | | | | | | | | | | | | | | | | | | | | | | | 0.441 | | | | | | | | 0.382 | | | | | | | | 0.0464 | 0.0989 | 63.7 | 23.6 | 12.7 |
| Hyper | | | | | | | | | | | | | | | | | | | | | | | | | | | | | | | | | | | | | | | | | - | - | - | - | - |
| Staring | | | | | | | | | | | | | | | | | | | | | | | | | | | | | | | | | | | | | | | | | - | - | - | - | - |
| SnapFlies | | | | | | | | | | | | | | | 3.280 | | | | | | | | | | | | | | | | | | | | | | | | | | 0.0105 | 0.0247 | 15.7 | 4.8 | 79.5 |
| TailChase | | | 1.003 | | | | | | | | | | | | | | 0.337 | | | 0.287 | | | | | | | | | | | | | | | | | | | | | 0.063 | 0.1353 | 73.5 | 25.1 | 1.4 |
| ShadowChase | | | | | | | | | | | | | | | | | | | | | | | | | | | | | | | | | | | | | | | | | - | - | - | - | - |
| Barking | | | | | | | | | | | | | | | | | | | | | | | | | | | | | | | | | | | | | | | | | - | - | - | - | - |
| GroomSelf | | 0.997 | | | | | | | | | | | | | | | | | | | | | | | | | | | | | | | | | | | | | | | 0.01 | 0.0262 | 48.5 | 43.9 | 7.6 |
| GroomOthers | 1.107 | | | | | | | | | | | | | | | | | | | | | | | | | | | | | | | | | | | | 0.314 | | | | 0.0576 | 0.1583 | 76.1 | 23 | 0.9 |
| OtherBehaviors | | | | | | | | | | | | | | | | | | | | | | | | | | | | | | | | | | | | | | | | | - | - | - | - | - |

Protective

Detrimental

**Supplementary Figure S11.** Logistic regression model selection using a stepwise selection method for traits recoded as case/control: case threshold at 90th percentile of the data. Odds ratio values are used to determine effect size and direction. Predicted probabilities and observed responses are displayed for each row.

95%

| | Weight | AcquirAGE | AgeAtEvaluation | Sex (Female vs. Male) | Neuter (Fixed vs. Intact) | Work (No vs. Yes) | Compete (No vs. Yes) | Purebred (No vs. Yes) | Pittbull (No vs. Yes) | AcquirPLACE (Breeder vs. Shelter) | AcquirPLACE (Other vs. Shelter) | AcquirPLACE (PetStore vs. Rescue) | AcquirPLACE (Rescue vs. Shelter) | OtherHOUSE (No vs. Yes) | Medication (No vs. Yes) | Medical (No vs. Yes) | Dogs (No vs. Yes) | Animals (No vs. Yes) | Kids (No vs. Yes) | Behavior (No vs. Yes) | Chr1A | Chr1B | Chr5 | Chr10A | Chr10B | Chr10C | Chr10D | Chr10E | Chr13 | Chr15A | Chr15B | Chr18 | Chr20 | Chr24A | Chr24B | Chr32 | Chr34 | ChrXA | ChrXB | ChrXC | R2 | Rescaled R2 | Concondant | Discordant | Tied |
|---|---|---|---|---|---|---|---|---|---|---|---|---|---|---|---|---|---|---|---|---|---|---|---|---|---|---|---|---|---|---|---|---|---|---|---|---|---|---|---|---|---|---|---|---|---|
| Train | | | | | | | 0.128 | | | | | | | | | 0.359 | | | | | | | | | | | | 7.407 | | | | | | | | 2.618 | | | | | 0.0662 | 0.2166 | 78.9 | 15.4 | 5.7 |
| StrDirAgg | | | | | | | | | | | | | | | | 0.113 | | | | 13.08 | | | | | | | | | | | | | | | | | | | | | 0.0682 | 0.2299 | 70.5 | 6.3 | 23.2 |
| OwnDirAgg | | | | | | | | | | | | | | | | | | | | | | | | | | | | | | | | | | | | | | | | | - | - | - | - | - |
| DogDirAgg | | | | | | | | | | | | | | | | | | | | | | | | | | | | | | | | | | | | | | | | | - | - | - | - | - |
| FamDogAgg | | | | | | | | | | | | | | | | | | | | | | | | | | | | | | | | | | | | | | | | | 0.1448 | 0.53 | 96.1 | 2.8 | 1.1 |
| DogDirFear | | | | | | | | | 2.977 | | | | | | | | | | | 4.597 | | | | | | | | | 2.879 | | | | | | | | | | | | 0.0482 | 0.1524 | 71 | 14.6 | 14.4 |
| StrDirFear | | | | | | | | | | | | | | | | | | | | | | | | | | | | | | | | | | | | | | | | | - | - | - | - | - |
| NonSocFear | | | | | | | | | | | | | | | | | | | | 6.513 | | | | | | | | | | | | | | | | | | | | | 0.036 | 0.1134 | 51.6 | 7.9 | 40.5 |
| TouchSen | | 0.997 | | | | | | | | | | | | | | | | | | 3.098 | 6.892 | | | | | | | | | | | | | | | | 9.028 | | | | 0.0681 | 0.2636 | 83.7 | 15.3 | 1 |
| SepRelProb | | 0.995 | 0.996 | | | 6.800 | | | | | | | | | | | | | | 6.236 | | | | | | 6.358 | | | 10.27 | | | | | | | | | | | | 0.0864 | 0.2832 | 84.9 | 15.1 | 0 |
| Excite | | | | | | | | | | | | | | | | | | | | | | | | | | | | | | | | | | | | | | | | | - | - | - | - | - |
| AtcAtnSeek | | | | | 8.333 | | | | | | | | | | | | | 0.352 | | | | | | | | | | | | | | | | | | | | | | | - | - | - | - | - |
| Chasing | | | | | | | | | | | | | | | | | | | | | | | | | | | | | | | | | | | | | | | | | - | - | - | - | - |
| Energy | | | | | | | | | | | | | | | | | | | | | | | | | | | | | | | | | | | | | | | | | - | - | - | - | - |
| EscapeRoam | | | | | | | | | | | | | | | | | | | | | | | | | | | | | | | | | | | | | | | | | - | - | - | - | - |
| Rolling | | | | | | | | | | | | | | | | | | | | | | | | | | | | | | | | | | | | | | | | | - | - | - | - | - |
| Coprophagia | | | | | | | | | | | | | | | | | | | | | | | | | | | | | | | | | | | | | | | | | - | - | - | - | - |
| Chewing | | | | | | | | | | | | | | | | | | | | | | | | | | | | | | | | | | | | | | | | | - | - | - | - | - |
| Mounting | | | | | | | | | | | | | | | | | | | | | | | | | | | | | | | | | | | | | | | | | - | - | - | - | - |
| Begging | | | | | | | | | | | | | | | | | | | | | | | | | | | | | | | | | | | | | | | | | - | - | - | - | - |
| FoodSteal | | | | | | | | | | | | | | | | | | | | | | | | | | | | | | | | | | | | | | | | | - | - | - | - | - |
| FearStairs | | | | | | | | | | | | | | | | | 0.216 | | | | | | | | | | | | | | | 4.193 | | | | 4.562 | 8.546 | | | | 0.0643 | 0.1953 | 75.8 | 19.8 | 4.4 |
| PullLeash | | | | | | | | | | | | | | | | | | | | | | | | | | | | | | | | | | | | | | | | | - | - | - | - | - |
| MarkUrine | | | | | | | | | | | | | | | | | | | | 9.416 | | | | | | | | | | | | | | | | | | | | | 0.0137 | 0.1058 | 56.1 | 6 | 37.9 |
| SubEmoUrn | | | | | | | | | | | | | | | | | | | | | | | | | | | | | | | | | | | | | | | | | - | - | - | - | - |
| SepUrn | | | | | | | | | | | | | | | | | | | | | | | | | | | | | | | | | | | | | | | | | - | - | - | - | - |
| SepDef | | | | | | | | | | | | | | | | | | | | 9.313 | | 0.046 | 151.5 | | | 6.896 | | | | | | 1.790 | 0.042 | | | | | | | | 0.1026 | 0.4794 | 96.1 | 3.6 | 0.3 |
| Hyper | | | | | | | | | | | | | | | | | | | | | | | | | | | | | | | | | | | | | | | | | - | - | - | - | - |
| Staring | | | | | | | | | | | | | | | | | | | | ##### | | | | | | | | | | | | | | | | | | | | | 0.0189 | 0.1262 | 58.4 | 5 | 36.6 |
| SnapFlies | | | | | | | | | | | | | | | | | | | 9.231 | 11.49 | | | | | | | | | | | | | | | | | | | | | 0.0494 | 0.2134 | 73.6 | 9.1 | 17.3 |
| TailChase | | | | | | | | | | | | | | | | | | | | | 4.155 | | | | | | | | | | | | | | | | | | | | 0.006 | 0.028 | 18.9 | 4.5 | 76.6 |
| ShadowChase | | | 1.006 | 0.347 | | | | | | | | | | | | | 0.289 | | | | | | | | | | | | | | | | 0.629 | | | | | | | | 0.0508 | 0.1767 | 79.5 | 19.9 | 0.5 |
| Barking | | | | | | | | | | | | | | | | | | | | | | | | | | | | | | | | | | | | | | | | | - | - | - | - | - |
| GroomSelf | | | | | | | | | | | | | | | | | | | | | | | | | | | | | | | | | | | | | | | | | - | - | - | - | - |
| GroomOthers | | | | | | | | | | | | | | | | | | | | | | | | | | | | | | | | | | | | | | | | | - | - | - | - | - |
| OtherBehaviors | | | | | | | | | | | | | | | | | | | | | | | | | | | | | | | | | | | | | | | | | - | - | - | - | - |

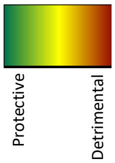


**Supplementary Figure S12.** Logistic regression model selection using a stepwise selection method for traits recoded as case/control: case threshold at 95th percentile of the data. Odds ratio values are used to determine effect size and direction. Predicted probabilities and observed responses are displayed for each row.

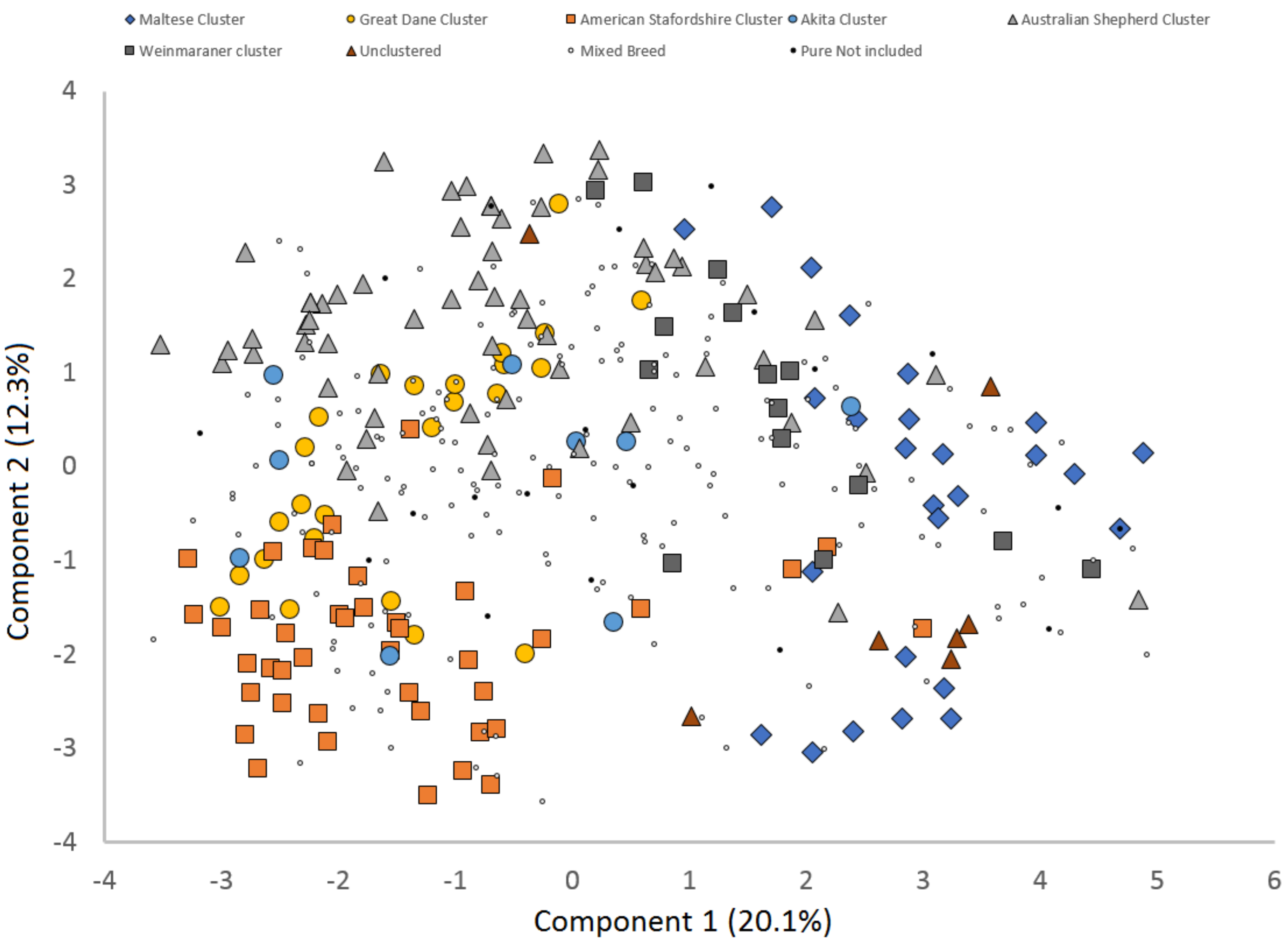


**Supplementary Figure S13.** Principal components analysis of genetic markers with classification of breeds according to Wilson et al. C-BARQ behavioral clustering (first two components). Note that the three AST cluster dogs >1 in PC1 are the three Italian Greyhounds, the only breed in the cluster that are not known to be related to all other members of this cluster. Technical note: Observations with missing values were omitted in the PCA, resulting in slightly different numbers of dogs in Figs. S9/S10. This is relevant for the PCA, but not for the modeling analyses.

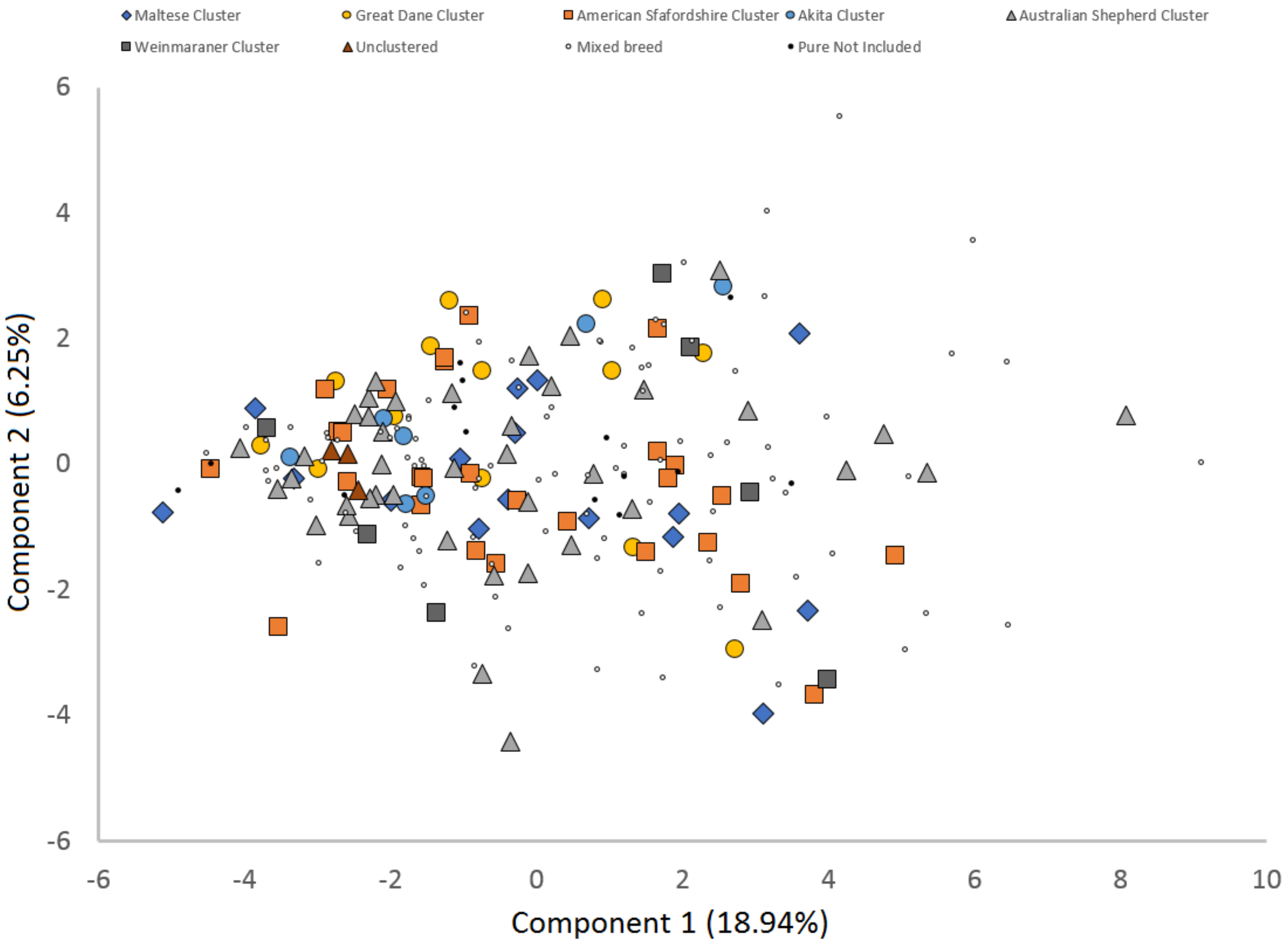


**Supplementary Figure S14.** Principal components analysis of C-BARQ data with classification of breeds according to Wilson et al. C-BARQ behavioral clustering (first two components). Technical note: Observations with missing values were omitted in the PCA, resulting in slightly different numbers of dogs in Figs. S9/S10. This is relevant for the PCA, but not for the modeling analyses.
